# A developmental analysis of juxtavascular microglia dynamics and interactions with the vasculature

**DOI:** 10.1101/2020.05.25.110908

**Authors:** Erica Mondo, Shannon C. Becker, Amanda G. Kautzman, Martina Schifferer, Christina E. Baer, Jiapei Chen, Eric J. Huang, Mikael Simons, Dorothy P. Schafer

## Abstract

Microglia, the resident macrophages of the central nervous system (CNS), are dynamic cells, constantly extending and retracting their processes as they contact and functionally regulate neurons and other glial cells. There is far less known about microglia-vascular interactions, particularly under healthy steady-state conditions. Here, we use the male and female mouse cerebral cortex to show that a higher percentage of microglia associate with the vasculature during the first week of postnatal development compared to older ages and the timing of these associations are dependent on the fractalkine receptor (CX3CR1). Similar developmental microglia-vascular associations were detected in the prenatal human brain. Using live imaging in mice, we found that juxtavascular microglia migrated when microglia are actively colonizing the cortex and became stationary by adulthood to occupy the same vascular space for nearly 2 months. Further, juxtavascular microglia at all ages contact vascular areas void of astrocyte endfeet and the developmental shift in microglial migratory behavior along vessels corresponded to when astrocyte endfeet more fully ensheath vessels. Together, our data provide a comprehensive assessment of microglia-vascular interactions. They support a mechanism by which microglia use the vasculature to migrate within the developing brain parenchyma. This migration becomes restricted upon the arrival of astrocyte endfeet when juxtavascular microglia then establish a long-term, stable contact with the vasculature.

**SIGNIFICANCE STATEMENT:** We report the first extensive analysis of juxtavascular microglia in the healthy, developing and adult brain. Live imaging revealed that juxtavascular microglia within the cortex are highly motile and migrate along vessels as they are colonizing cortical regions. Using confocal, expansion, super-resolution, and electron microscopy, we determined that microglia associate with the vasculature at all ages in areas lacking full coverage astrocyte endfoot coverage and motility of juxtavascular microglia ceases as astrocyte endfeet more fully ensheath the vasculature. Our data lay the fundamental groundwork to investigate microglia-astrocyte crosstalk and juxtavascular microglial function in the healthy and diseased brain. They further provide a potential vascular-dependent mechanism by which microglia colonize the brain to later regulate neural circuit development.

## INTRODUCTION

While myeloid lineage in origin, microglia are now appreciated to be key cellular components of neural circuits. Imaging studies have revealed that microglia are constantly extending and retracting their processes, which are in frequent contact with neurons, synapses, and other glial cells (Davalos et al. 2005; Nimmerjahn, Kirchhoff, and Helmchen 2005; Schafer et al. 2012; Tremblay, Lowery, and Majewska 2010; Frost and Schafer 2016). These descriptions of physical interactions between microglia and other resident CNS cell types have now led to a new understanding that microglia are important for neural circuit structure and function, including their role in developmental synaptic pruning by engulfing and removing synapses from less active neurons (Schafer et al. 2012; Tremblay, Lowery, and Majewska 2010; Paolicelli et al. 2011; Gunner et al. 2019). Besides interactions with parenchymal neurons and glia, microglia are known to interact with the vasculature. However, the vast majority of these studies have been in the context of disease where parenchymal microglia rapidly associate with the brain vasculature following breakdown of the blood-brain barrier (BBB) and, in turn, inflammatory microglia can modulate the breakdown of the BBB (Stankovic, Teodorczyk, and Ploen 2016; Zhao et al. 2018). Far less is known about how microglia interact with the vasculature in the healthy brain. With new evidence that microglia could be a conduit by which changes in peripheral immunity (e.g. microbiome, infection, etc.) affect CNS function (Hanamsagar and Bilbo 2017; Hammond, Robinton, and Stevens 2018; Zhao et al. 2018; Rothhammer et al. 2018) and mounting evidence that an array of neurological disorders have a vascular and microglial component (Daneman 2012; Hammond, Robinton, and Stevens 2018; Zhao et al. 2018), a greater understanding of microglia-vascular interactions is necessary.

The neurovascular unit (NVU) is composed of endothelial cells, pericytes, vascular smooth muscle cells, astrocytes, macrophages, and neurons that connect the brain parenchyma to the cerebral vasculature. Interactions between these NVU cell types is important for a variety of physiological processes such as angiogenesis, vessel maintenance and permeability, metabolic support, and regulation of blood flow (Brown et al. 2019; McConnell et al. 2017). The development of the NVU begins around embryonic day (E) 9.5 in mice, when specialized endothelial cells branch from vessels of the perineural vascular plexus to form capillaries that invade nearby neural tissue (Saili et al. 2017). Pericytes associate with endothelial cells as nascent vessels generate at E9.5 (Armulik et al. 2010; Bauer et al. 1993; Yamanishi et al. 2012; Daneman et al. 2010) and these interactions are critical to form the BBB (Zlokovic 2008; Daneman et al. 2010). Astrocytes are also a key component of the mature NVU. After the vasculature initially forms, astrocytes extend their processes to form endfeet over the course of postnatal development in rodents (Daneman et al. 2010). These astrocyte endfeet ultimately surround and ensheath the majority of the vasculature by adulthood where they play roles in a variety of functions such as maintaining the BBB, providing metabolic support to neurons, and regulating blood flow (Abbott, Rönnbäck, and Hansson 2006; Kimelberg and Nedergaard 2010; Macvicar and Newman 2015).

The vast majority of studies assessing interactions between microglia and the vasculature are in the context of disease. For example, microglia rapidly surround and contact the vasculature following breakdown of the BBB in the inflamed CNS (Zhao et al. 2018; Stankovic, Teodorczyk, and Ploen 2016). One mechanism regulating these microglia-vascular interactions is the blood component fibrinogen and CD11b on microglia (Davalos et al. 2012; Adams et al. 2007). Reactive microglia can also influence the opening of the BBB by phagocytosing astrocyte endfeet or upregulating molecules such as VEGF, iNOS, and ROS (Stankovic, Teodorczyk, and Ploen 2016; Zhao et al. 2018; Haruwaka et al. 2019). In the healthy brain, much less is known. Studies in rodents and humans have shown that microglia associate with the vasculature in the developing CNS and live imaging in postnatal brain slices following traumatic injury or in embryonic mouse brain slices has suggested that microglia can migrate along the vasculature (Monier et al. 2007; Fantin et al. 2010; Smolders et al. 2017; Grossmann et al. 2002; Checchin et al. 2006). Microglia have also been suggested to regulate vascular growth and complexity in the developing hindbrain and retina (Fantin et al. 2010; Rymo et al. 2011; Checchin et al. 2006; Yoshiaki Kubota et al. 2009; Dudiki et al. 2020). Together, these studies provide evidence that there is microglia-vascular crosstalk, which requires further investigation in development, adulthood, and disease.

In the current study, we investigated microglia-vascular interactions in the healthy, developing and adult cerebral cortex. Using confocal, super-resolution, expansion, and electron microscopy, we assessed the developmental regulation of physical associations between microglia and the vasculature and used fractalkine receptor (CX3CR1)-deficient mice to determine a role for this signaling in these physical associations. Using *in vitro* confocal and *in vivo* 2-photon live imaging, we further assessed the dynamics of juxtavascular microglia in real time. Our data support a mechanism by which microglia migrate along the vasculature to colonize the developing brain and the timing of these interactions is regulated by CX3CR1. This migratory behavior becomes restricted as astrocyte endfeet mature and suggests the establishment of a long-term niche for juxtavascular microglia in the adult brain.

## MATERIALS AND METHODS

### Animals

Male and female mice were used for all experiments. *Cx3cr1^-/-^* mice (Cx3cr1^EGFP/EGFP^; stock #005582) and *C57Bl6/J* (stock #000664) mice were obtained from Jackson Laboratories (Bar Harbor, ME). Heterozygous breeder pairs were set up for all experiments and wild-type (WT) and heterozygote littermates were used as controls with equal representation of males and females for each genotype. All experiments were performed in accordance with animal care and use committees and under NIH guidelines for proper animal welfare.

### Human prenatal brain collection and immunofluorescence microscopy

Deidentified prenatal human brain tissues were collected via the Department of Pathology Autopsy Service at the University of California San Francisco under the approval of the Committee on Human Research (CHR, Study #: 12-08643). Brain tissues from four prenatal cases at 15, 18, 21 and 28 gestational weeks (GW) were evaluated using standard neuropathologic examinations to rule out any gross or microscopic abnormalities. These autopsy cases, which all had postmortem intervals of less than 48 hours, were fixed in freshly prepared 4% paraformaldehyde (PFA) and sampled at the level of the mammillary body. Following fixation in 4% PFA for 48 hours, brain samples were incubated with 20% sucrose solution, and were frozen in embedding medium OCT for cryosectioning at 20µm. For consistency, 3-6 consecutive sections were prepared from each sample and immunostained with anti-Iba1 antibody (Wako; Richmond, VA; 1:3000) and anti-CD31 antibody (R&D Systems; Minneapolis, MN; 1:200). Images of the ventricular and subventricular zones at the level of the frontal cortex were acquired on Leica SP8 confocal microscope using a 40X (1.3NA) objective lens.

### Preparation of tissue for immunofluorescence microscopy

Mice were perfused with 1X Hank’s balanced salt solution (HBSS) -magnesium, -calcium, (Gibco, Gaithersburg, MD) prior to brain removal at indicated ages. For analysis of frontal and somatosensory cortex, brains were post-fixed in 4% paraformaldehyde in 0.1M phosphate buffer (PB) for four hours. Brains were placed in 30% sucrose in 0.1M PB and allowed to sink prior to sectioning. Sections were blocked in 10% goat serum, 0.01% TritonX-100 in 0.1M PB for 1 hour before primary immunostaining antibodies were applied overnight. Secondary antibodies were applied for two hours the following day. All steps were carried out at room temperature with agitation. For structured illumination microscopy (SIM), sections were blocked in 3% BSA, 0.01% TritonX-100 in 0.1M PB for 1 hour before primary immunostaining antibodies were applied for 48 hours at 4°C. Secondary antibodies were applied for four hours at room temperature with agitation. The following antibodies were used: anti-P2RY12 (Butovsky Laboratory, Brigham and Women’s Hospital, Harvard University; 1:200), anti-PECAM (Biolegend; San Diego, CA; 1:100), anti-aquaporin 4 (Millipore Sigma; St. Louis, Missouri; 1:200), anti-Pdgfrβ (Thermo Fisher Scientific; Waltham, MA; 1:200), anti-Lyve1 (Abcam; Cambridge, MA; 1:200), anti-smooth muscle actin (SMA) (Millipore Sigma; St. Louis, Missouri; 1:200) and anti-VGluT2 (Millipore Sigma; St. Louis, Missouri; 1:2000).

### Confocal microscopy

Immunostained sections were imaged on a Zeiss Observer Spinning Disk Confocal microscope equipped with diode lasers (405nm, 488nm, 594nm, 647nm) and Zen acquisition software (Zeiss; Oberkochen, Germany). For microglia-vascular interaction, microglial density, microglia association with SMA+ or SMA- vessels and vascular density analyses, 20x, single optical plane, tiled images of the frontal or somatosensory cortex were acquired for each animal. To create a field of view (FOV), each tiled image was stitched using Zen acquisition software. Two FOVs (ie. tiled images) were acquired per animal. To note, anti-P2RY12 immunostaining was used to label microglia in wild type animals, which was more difficult to visualize at lower magnification at older ages compared to EGFP-labeled microglia. As a result, for anti-P2RY12 immunostained sections from P7-P28 animals, twelve 40x fields of view were acquired per animal with 76 z-stack steps at 0.22µm spacing. For analysis of vascular diameter, juxtavascular association with branched/unsegmented vessels, primary processes aligned with vessels, astrocyte endfeet coverage on the vasculature, and vascular-associated microglia contacts with astrocytes, six-twelve 40X fields of view were acquired from the frontal cortex per animal with 76 z-stacks at 0.22µm spacing.

### Juxtavascular microglia and microglia density analyses in the frontal and somatosensory cortices

Using the DAPI channel as a guide, a region of interest (ROI) was chosen in each cortical layer, I-VI from each 20x stitched tiled image (10 ROIs per animal). Subsequent images were analyzed in ImageJ (NIH; Bethesda, MD). For anti-P2RY12, sections were acquired at 40x, a maximum intensity projection was made from each z-stack and was considered a ROI (12 per animal). The ROI areas were recorded. The same ROI was transposed on the microglial channel and the cell counter ImageJ plugin was used to count the number of microglia in the ROI. The total density of microglia was then calculated by dividing the microglia number by the ROI area. To assess microglial association with the vasculature, the microglia and blood vessel channels were merged and the cell counter plugin was used to manually count the number of microglia with cell bodies contacting blood vessels. Juxtavascular microglia were defined as microglia with at least 30% of their soma perimeter in contact with blood vessels and soma centers that were within 10µm of the vessel. The percent of juxtavascular microglia was calculated by summing the total number of microglia on vasculature divided by the total number of microglia within the ROI. For each animal, data from the ROIs were averaged together to get a single average per animal for statistical analyses.

### Juxtavascular microglia analysis within the barrel cortex

Juxtavascular microglia analysis in the barrel cortex was performed blinded to genotype. Images were analyzed in ImageJ (NIH; Bethesda, MD). From each tiled image from each animal, 12-18 images containing VGluT2+ barrels were cropped for subsequent analyses. From each cropped image, the individual channels were separated and, using the free hand selection tool, each individual barrel was outlined. This ROI outlining the barrel was transposed to the microglia channel where the cell counter plugin was used to count the number of microglia in the barrels. The microglia and blood vessel channels were then merged and the same ROI was transposed onto the merged image. The cell counter plugin was used to count the number of microglia in barrels associated with vasculature. Each individual barrel ROI was then cleared, leaving behind only the septa fluorescence and the cell counter plugin was again used to count the number of microglia and the number of juxtavascular microglia in the septa. To calculate the percent of juxtavascular microglia in the barrel cortex, the total numbers of juxtavascular microglia in the barrels and septa were summed and divided by the total number of microglia in the barrel and septa, respectively, for each ROI. The total microglia in barrels and septa, regardless of vascular association, were also calculated. All numbers across 12-18 cropped images were then averaged for a given animal prior to statistical analyses.

### Vascular density analysis

Density analysis was performed blinded to genotype from the same tiled and stitched 20x images used for microglia-vascular association analyses. Using ImageJ (NIH; Bethesda, MD) software, the blood vessel channel was thresholded manually and the total blood vessel area was measured. Vascular density was calculated by dividing the blood vessel area by the area of the ROI. For each animal, the vascular density was averaged across all ROIs in the two FOV to get a single average per animal for statistical analyses.

### Microglial association with SMA+ or SMA- vessels analysis

Using the DAPI channel as a guide, a region of interest (ROI) was chosen in each cortical layer, I-VI from each 20x stitched tiled image (10 ROIs per animal). Subsequent images were analyzed in ImageJ (NIH; Bethesda, MD). The same ROI was transposed on the microglial, Pdgfrβ, and SMA channel and the cell counter ImageJ plugin was used to count the total number of microglia, the number of juxtavascular microglia, and the number of juxtavascular microglia contacting SMA+ or SMA- vessels in the ROI. The percent of juxtavascular microglia contacting SMA+ or SMA- vessels was quantified by dividing the number of microglia on SMA+ or SMA- vessels by the number of total juxtavascular microglia. For each animal, data from the ROIs were averaged together to get a single average per animal for statistical analyses.

### Vascular diameter analysis

Using Imaris (Bitplane) software, the diameter of the vessel was measured in 3D at microglial soma contact points from 40X images (12 per animal). For each animal, data from the 12 images were averaged together to get a single average per animal for statistical analysis.

### Primary Process and branched/unsegmented vessel analyses

Using ImageJ (NIH; Bethesda, MD), the total number of primary processes, the number of primary processes aligned parallel with vessels, and whether the juxtavascular microglia was contacting a vessel branch point was calculated from 40X images (6 per animal, n=3-4 animals). The percent of primary processes aligned with vessels was calculated by dividing the number of primary processes aligned parallel and in direct contact with vessels by the total number of primary processes. The percent of juxtavascular microglia contacting branched/unsegmented vessel was calculated by dividing the number of juxtavascular microglia contacting branched or unsegment vessels by the total number of juxtavascular microglia. For each animal, data from 6 images were averaged together to get a single average per animal for statistical analysis.

### Acute Slice Time-Lapse Imaging

Mice were given a retro-orbital injection of Texas Red labeled dextran (Fisher Scientific; Waltham, MA) 10 minutes prior to sacrifice to label vasculature. Mice were euthanized at P7 or P≥120, brains were isolated and sectioned coronally at a thickness of 300µm using a Leica VT1200 vibratome in oxygenated 37°C artificial cerebrospinal fluid (ACSF). Slices were mounted on a MatTak glass bottom microwell dish and placed in a Zeiss Observer Spinning Disk Confocal microscope equipped with diode lasers (405nm, 488nm, 594nm, 647nm) and Zen acquisition software (Zeiss; Oberkochen, Germany). Image acquisition started after a minimum of 30 minutes of tissue equilibration at 37°C with 5% CO_2_ and within 2 hours of decapitation. Oxygenated ACSF was continuously perfused over the slices at a rate of 1.5-2µm/minute for the duration of equilibration and imaging. Per animal, one field of view was imaged every 5 minutes over 6 hours on an inverted Zeiss Observer Spinning Disk Confocal and a 20X objective. Z-stacks spanning 50-60µm, with serial optical sections of 1.5-2µm were recorded from a minimal depth of 30µm beneath the surface of the slice to avoid cells activated by slicing.

### *In vivo* 2-Photon Time-Lapse Imaging

Cranial window surgeries were performed as previously described within the visual cortex (2.5µm lateral and 2.0 µm posterior from bregma) (Goldey et al. 2014). One week after surgery, mice were head-fixed to a custom-built running wheel and trained to run while head restrained for increasing time intervals several days a week. Two weeks post surgery long-term 2-photon live imaging began. Mice were given a retro-orbital injection of Texas Red labeled dextran (Fisher Scientific; Waltham, MA) 10 minutes prior to imaging and were head restrained on a custom built running wheel, which was positioned directly under the microscope objective. Images were acquired with a 20X water immersion objective (Zeiss, NA 1.0) on a Zeiss Laser Scanning 7 MP microscope equipped with a tunable coherent Chameleon Ultra II multiphoton laser and BiG detector. Three different regions of interest (ROIs) were taken at least 75µm below the surface of the brain, with z-stacks spanning 45-65µm with a step size of 2.5µm for each animal. On the first day of imaging, each ROI was imaged every 5 minutes over 2 hours. The same ROIs were then imaged once (single z-stack) on the following days post first imaging session: 1, 3, 7, 10, 14, 17, 21, 24, 28, 35, and 42 days. For each imaging day, the ROIs from day 0 of imaging were identified based on the vascular structure.

### Migration tracking and analysis

Image processing and microglial soma motility/migration tracking were performed using ImageJ (NIH; Bethesda, MD). Time series were first corrected for 3D drift using the 3D drift correction plugin (Parslow, Cardona, and Bryson-richardson 2014) and migration was tracked using the TrackMate plugin (Tinevez et al. 2017). For each developmental time point, 10-12 juxtavascular and vascular-unassociated microglia were analyzed per animal (n=4 mice per developmental time point). Only cells remaining in the field of view for six hours were included in the analysis. The average soma motility (µm/h) was calculated by measuring the displaced distance of the microglial soma between time=0 min and time=360 min and dividing by the duration of the imaging session. Juxtavascular distance migrated was calculated by measuring the displaced distance of the microglial soma between time=0 min and time=360 min. Juxtavascular migration trajectory was calculated by measuring the angle between the blood vessel and juxtavascular microglia soma along the longest, continuous stretch of motility on the vessel. Percent of cells within each binned category (motility, distance travelled, and trajectory) was calculated by dividing the number microglia of within each category by the total number of microglia. For each animal, data from each analyzed cell were averaged together to get a single average per animal for statistical analysis.

*In vivo* tracking of juxtavascular microglia motility and long-term juxtavascular microglia were performed using ImageJ (NIH; Bethesda, MD). Time series were first corrected for 3D drift using the 3D drift correction plugin (Parslow, Cardona, and Bryson-richardson 2014) and migration was tracked using the TrackMate plugin (Tinevez et al. 2017). To calculate percent of microglia stationary over two hours, the number of stationary juxtavascular and vascular-unassociated microglia was divided by the total number of microglia. To calculate the percent of original juxtavascular microglia that remain on vessels over 42 days, the number of juxtavascular microglia on day 0 was calculated. For each subsequent day, the number of these original juxtavascular microglia that were still associated with the vasculature was determined and divided by the number of original juxtavascular microglia on day 0. For each animal, data was analyzed from three ROIs and averaged together to get a single average per animal for statistical analysis. For each animal, data was analyzed from three ROIs and added together to get a single average per animal for statistical analysis.

### Astrocyte endfeet coverage analysis

Using Imaris (Bitplane) software, the astrocyte endfeet and vessel channels were 3D rendered from 40X images (6 per animal). The astrocyte channel was then masked onto the vessel channel and the masked astrocyte channel was 3D rendered. Volumes of the 3D rendered vessel channel and masked astrocyte endfeet channel were recorded. The percent of blood vessels covered by astrocyte endfeet was calculated by dividing the blood vessel volume by the masked astrocyte endfeet volume. For each animal, data from the 6 images were averaged together to get a single average per animal for statistical analysis.

### Juxtavascular microglia-astrocyte contact

Analysis was done using the same images used for astrocyte endfeet coverage analysis in Imaris (bitplane). The microglia was 3D rendered, masked onto the blood vessel and astrocyte endfeet channel, and the volume of the masked microglial channel was recorded. The percent of juxtavascular microglia contacting blood vessels only, vessels and astrocyte endfeet, or astrocyte endfeet only was calculated by summing the number of microglia contacting vessels only, vessels and astrocyte endfeet, or astrocyte endfeet only and dividing by the total number of juxtavascular microglia. For each animal, data from the 6 images were averaged together to get a single average per animal for statistical analysis.

### Expansion Microscopy (ExM)

Expansion microscopy was performed as previously described (Asano et al. 2018) with slight modification. Briefly, 80µm floating sections were blocked in 0.5% bovine serum albumin (BSA) and 0.3% Triton-X100 (TX-100) for 1 hour at room temperature. Primary antibodies, anti-aquaporin 4 (Millipore Sigma; St. Louis, Missouri; 1:200), anti-PDGFRβ (Thermo Fisher Scientific; Waltham, MA; 1:100), and anti-GFP (Abcam; Cambridge, MA; 1:200) were incubated in 0.5% BSA and 0.3% TX-100 at 4°C for 4 nights. Secondary antibodies were added at 1:200 dilutions overnight at room temperature. Expansion microscopy protocol (Basic Protocol 2) was then followed as published in Asano et al. 2018.

### Structured Illumination Microscopy (SIM)

Structured Illumination Microscopy (SIM) was performed using a GE Delta Vision OMX V4 microscope with pCO.edge sCMOS cameras and an Olympus 60x 1.42 NA objective. Samples were mounted in Prolong Glass mounting media with #1.5 coverslips and imaged using 1.516 refractive index immersion oil. Image processing was completed using the GE softWorx software and image quality was determined using the SIMcheck plugin in ImageJ. SIM figures were produced in ImageJ (NIH; Bethesda, MD).

### Scanning Electron Microscopy (SEM)

Mice were perfusion fixed in 2.5% glutaraldehyde and 2% paraformaldehyde in 0.1 M sodium cacodylate buffer at pH 7.4 (Science Services). Brains were dissected, vibratome sectioned, and immersion fixed for 24h at 4°C. We applied a rOTO (reduced osmium-thiocarbohydrazide-somium) staining procedure adopted from Tapia et al. (Tapia et al. 2013). Briefly, the tissue was washed and post-fixed in 2% osmium tetroxide (EMS), 2% potassium hexacyanoferrate (Sigma) in 0.1 M sodium cacodylate buffer. After washes in buffer and water the staining was enhanced by reaction with 1% thiocarbohydrazide (Sigma) for 45 min at 50°C. The tissue was washed in water and incubated in 2% aqueous osmium tetroxide. All osmium incubation steps were carried out over 90 min with substitution by fresh reagents after 45 min, respectively. To further intensify the staining, 2% aqueous uranyl acetate was applied overnight at 4°C and subsequently warmed to 50°C for 2h. The samples were dehydrated in an ascending ethanol series and infiltrated with LX112 (LADD). The samples were flat embedded into gelatin capsules (Science Services) and cured for 48h. The block was trimmed by 200 µm at a 90° angle on each side using a TRIM90 diamond knife (Diatome) on an ATUMtome (Powertome, RMC). Consecutive sections were taken using a 35° ultra-diamond knife (Diatome) at a nominal cutting thickness of 100 nm and collected on freshly plasma-treated (custom-built, based on Pelco easiGlow, adopted from Mark Terasaki) CNT tape (Yoshiyuki Kubota et al. 2018). We collected 450 (P5) and 550 (P56) cortical sections, covering a thickness of 45-55 µm in depth. Tape strips were mounted with adhesive carbon tape (Science Services) onto 4-inch silicon wafers (Siegert Wafer) and grounded by additional adhesive carbon tape strips (Science Services). EM micrographs were acquired on a Crossbeam Gemini 340 SEM (Zeiss) with a four-quadrant backscatter detector at 8 kV. In ATLAS5 Array Tomography (Fibics), the whole wafer area was scanned at 3000 nm/pixel to generate an overview map. The entire ultrathin section areas of one wafer (314 sections (P5), 279 sections (P56)) were scanned at 100 x 100 x 100 nm^3^ (465 x 638 µm^2^ (P5), 1249 x 707 µm^2^ (P56). After alignment in Fiji TrakEM2 (Cardona et al. 2012) areas that contained microglia in close proximity to blood vessels (148 x 136 x 16 µm^3^ (P5), 193 x 186 x 12 µm^3^ (P56) were selected for high resolution acquisition. We collected 29 total 2D micrographs (10 x 10 nm^2^) from n=3 animals at P5 and 11 total micrographs from n=3 animals at P56. From each age, one juxtavascular microglia was identified and selected to generate a 3D volume (10 x 10 x100 nm^3^). The image series were aligned in TrakEM2 using a series of automated and manual processing steps. For the P5 and P56 image series, segmentation and rendering was performed in VAST (Volume And Segmentation Tool) (Berger et al. 2018). We used Blender to render the two 3D models (Community 2018).

### Statistical analyses

GraphPad Prism 7 (La Jolla, CA) provided the platform for all statistical and graphical analyses. The ESD method was run for each ROI per animal to identify outliers. Significant outliers were removed prior to analyses. Analyses included Students t-test when comparing 2 conditions or one-way ANOVA followed by Dunnett’s post hoc analysis or two-way ANOVA followed by Sidak’s or Tukey’s post hoc analyses (indicated in figure legends).

## RESULTS

### A high percentage of microglia are juxtavascular during development

During rodent and human embryonic development, microglia somas have been described to be in close contact with blood vessels (Fantin et al. 2010; Monier et al. 2007; Checchin et al. 2006). We assessed microglial association with the vasculature over an extended developmental time course across postnatal development. Microglia were labeled using transgenic mice that express EGFP under the control of the fractalkine receptor CX3CR1 (*Cx3cr1^EGFP/+^*). The vasculature was labeled with an antibody against platelet endothelial cell adhesion molecule (PECAM). To start, we focused our analyses in the frontal cortex. Juxtavascular microglia were defined as microglia with at least 30% of their soma perimeter in contact with blood vessels and soma centers that were within 10µm of the vessel, which we confirmed with orthogonal views and 3D surface rendering (Fig 1. A-F; See also Movies 1 and 2). Juxtavascular microglia were further distinguished from perivascular macrophages by their morphology with processes emanating from their soma and higher levels of EGFP. Using these criteria, we found a higher percent of the total microglial population were juxtavascular at P1-P5 (Fig. 1G) in the frontal cortex, which was independent of sex (data not shown). The percent association dropped to below 20% by P14 and was maintained at later ages. We confirmed that this developmental regulation of juxtavascular microglia was independent of changes in vasculature density over development. While the total vascular content of the cortex increases as the brain grows, the density of the blood vessels within a given field of view is unchanged across development (Fig. 1H). Consistent with the results in mouse, the ventricular and subventricular zones of the prenatal human brain at the level of the frontal cortex also showed a high percent of juxtavascular microglia. This association in the developing human brain peaked at 18-24 gestational weeks (GW) where 38% of total microglia were juxtavascular (Fig. 1I-J)—a percentage similar to what we identified in early postnatal mice. Together, these data demonstrate that a large percentage of the total microglia are juxtavascular in the early postnatal mouse and prenatal human brain.

**Figure 1:**
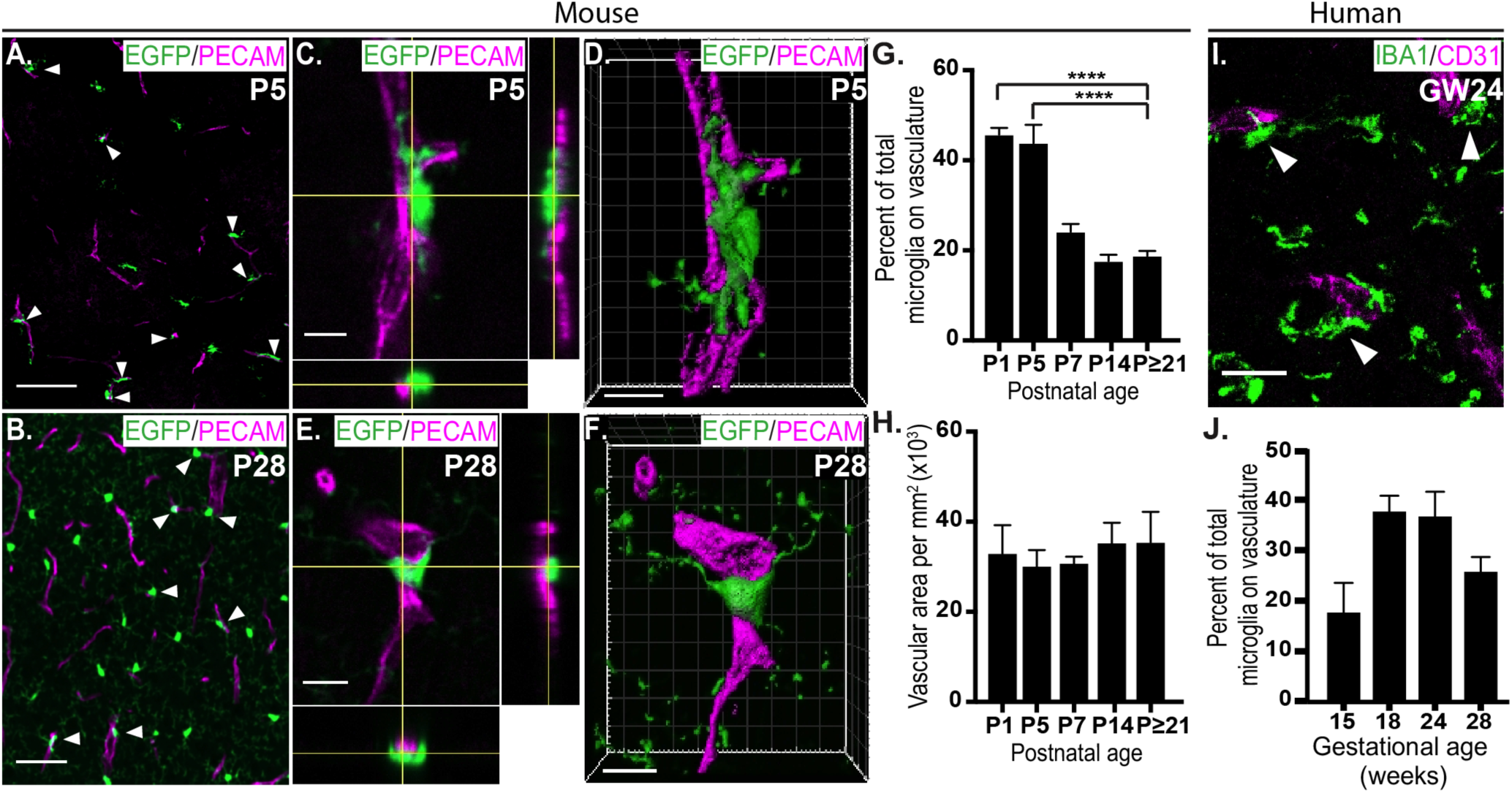
A high percentage of microglia are juxtavascular during early postnatal development. **A-B.** Representative low magnification tiled images of microglia (green, EGFP) associated with vasculature (magenta, anti-PECAM) in the P5 (**A**) and P28 (**B**) frontal cortex. Filled arrowheads denote juxtavascular microglia. Scale bars= 100 µm (**A**) and 50 µm (**B**). **C-D.** High magnification, orthogonal view (**C**) and 3D reconstruction and surface rendering (**D**) of juxtavascular microglia in the P5 frontal cortex (see also Movie 1). Scale bars= 10 µm. **E-F**. Orthogonal (**E**) and 3D reconstruction and surface rendering (**F**) of a juxtavascular microglia in the P28 frontal cortex (see also Movie 2). Scale bars= 10 µm. **G.** The percent of the total microglia population associated with vasculature over development in the frontal cortex. One-way ANOVA with Dunnett’s post hoc; comparison to P≥21, n=4 littermates per developmental time point, ****p<.0001. **H**. Vascular density over development in the frontal cortex. One-way ANOVA with Dunnett’s post hoc; comparison to P≥21, n=4 littermates per developmental time point. **I**. Representative image of microglia (green, anti-IBA1) associated with vasculature (magenta, anti-CD31) in gestational week (GW) 24 in the ventricular zone (VZ) and subventricular zone (SVZ) at the level of the human frontal cortex. Filled arrowheads denote juxtavascular microglia. Scale bar**=**20 µm. **J.** Quantification of the percentage of total microglia associated with vasculature in the human brain. One-way ANOVA across all ages, p=0.0544, n=1 specimen per gestational age. All error bars represent ± SEM.

### Juxtavascular microglia are largely associated with capillaries in the early postnatal cortex

While previous work has described similar high association of microglia with the vasculature in the embryonic/prenatal brain, these studies did not use markers to distinguish microglia from perivascular macrophages (Fantin et al. 2010; Monier et al. 2007; Checchin et al. 2006). Therefore, we next sought to confirm that vascular-associated EGFP-positive cells were, indeed, microglia versus perivascular macrophages and determine which types of vessels were being contacted by microglia. We found that the juxtavascular EGFP+ cells that we initially identified as microglia based on their larger numbers of processes and higher levels of EGFP (Fig. 1; Fig. 2A-B filled arrowheads) were also positive for the microglia-specific marker P2RY12 (Fig. 2A, filled arrowhead) and negative for the perivascular macrophage-specific marker LYVE1 (Fig. 2B, unfilled arrowheads) (Butovsky et al. 2014; Zeisel et al. 2015). Using anti-P2RY12 to label microglia in wild-type mice or EGFP in *Cx3cr1^EGFP/+^* mice, which are heterozygote for CX3CR1, we obtained similar percentages of juxtavascular microglial and vascular density (Fig. 2C-D), confirming results were independent of the microglial labeling technique. We also found that these juxtavascular microglia were associated largely along unsegmented vessels, rather than branch points, across postnatal development (Fig. 2E). We next assessed what types of vessels were contacted by juxtavascular microglia, using a combination of parameters. Capillaries are ≤8 µm in diameter and are smooth muscle actin (SMA)-negative and Platelet Derived Growth Factor Receptor β (PDGFRβ)-positive (Grant et al. 2019; Mastorakos and Mcgavern 2019). Arterioles are >8 µm in diameter and are SMA-positive and a subset of pre-capillary arterioles are also PDGFRβ-positive (Grant et al. 2019). Using these markers, we identified that juxtavascular microglia were largely contacting capillaries (≤8 µm, SMA-negative, PDGFRβ-positive; Fig. 2F-H). These experiments establish that a large percentage of bona fide microglia are associated with unsegmented capillaries in the postnatal cerebral cortex and these percentages are similar in wild type and *Cx3cr1^EGFP/+^* mice.

**Figure 2:**
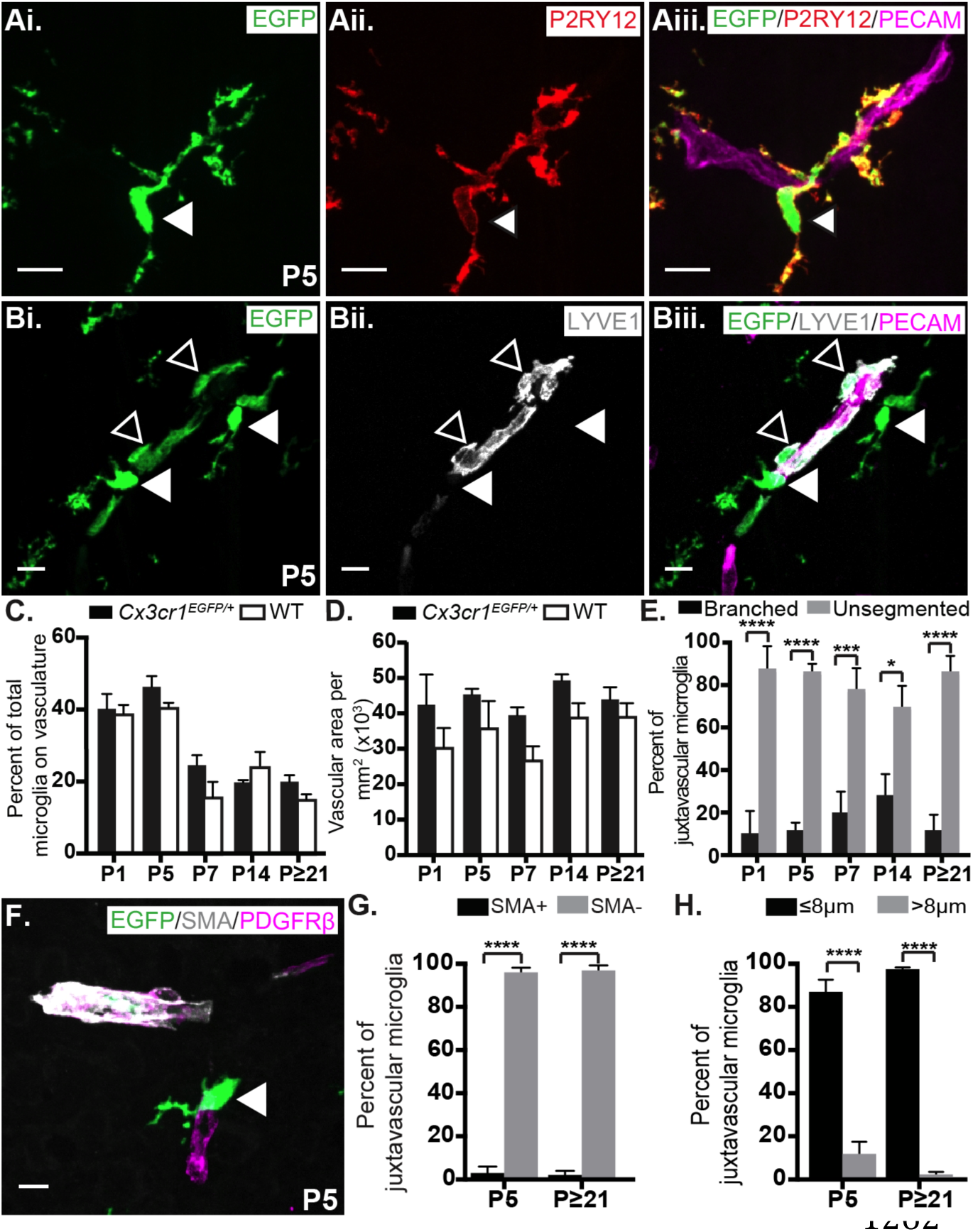
Juxtavascular microglia predominantly contact capillaries in the postnatal cortex. **A.** A representative image of a juxtavascular microglia (filled arrowhead) in the P5 frontal cortex. Microglia are labeled using the *Cx3cr1^EGFP/+^* reporter mouse (green; **Ai**) and immunolabeling for a microglia-specific marker anti-P2RY12 (red; **Aii**.). The vasculature is labeled with anti-PECAM (magenta) in the merged image (**Aiii.**). Scale bar= 10µm. **B.** A representative image of LYVE1-negative microglia (green, EGFP, filled arrowheads) and LYVE1-positive perivascular macrophages (gray, anti-LYVE1, unfilled arrowheads) associated with vasculature (magenta, anti-PECAM) in the P5 frontal cortex. Scale bar= 10µm. **C.** Quantification of juxtavascular microglia across development labeled either with EGFP in *Cx3cr1 ^EGFP/+^* mice (black bars) or anti-P2RY12 in wild type mice (WT, white bars) frontal cortices. Two-way ANOVA with a Sidak’s post hoc; n=3-4 littermates per genotype per developmental time point. **D**. Quantification of vascular density in *Cx3cr1 ^EGFP/+^* (black bars) and WT (white bars) frontal cortices over development. Two-way ANOVA with a Sidak’s post hoc; n=3-4 littermates per genotype per developmental time point. **E.** Quantification of the percent of juxtavascular microglia contacting branched (black bars) or unsegmented (gray bars) vessels. Two-way ANOVA with a Sidak’s post hoc; n=3-4 littermates per developmental time point, *p<.05, ***p<.001, ****p<.0001. **F.** A representative image of a juxtavascular microglia (green, EGFP, filled arrowhead) contacting smooth muscle cell actin (gray, SMA)-negative capillaries (magenta; PDGFRβ) in the P5 frontal cortex. Scale bar= 10µm **G.** Quantification of the percent of juxtavascular microglia contacting SMA-positive or -negative vessels at P5 and P≥21 in the frontal cortex. Two-way ANOVA with a Sidak’s post hoc; n=3 littermates per genotype per developmental time point, ****p<.0001. **H.** Quantification of the percent of juxtavascular microglia contacting vessels ≤8µm and >8µm at P5 and P≥21 in the frontal cortex. Two-way ANOVA with a Sidak’s post hoc; n=4 littermates per genotype per developmental time point, ****p<.0001. All error bars represent ± SEM.

### High percentages of juxtavascular microglia occur when microglia are actively colonizing the cortex in a CX3CR1-dependent manner

Over development, microglia undergo a dynamic process of colonization and expansion in a rostral-to-caudal gradient (Ashwell 1991; Perry, Hume, and Gordon 1985). Similar to previously published work (Nikodemova et al. 2015), we identified a large expansion in cortical microglia between P1 and P14, with microglia colonizing the more rostral frontal cortex region prior to the more caudal somatosensory cortex (Fig. 3A-C, bar graphs in B-C). Microglia-vascular association mirrored this rostral-to-caudal gradient by which microglia colonize the brain with a higher percentage of juxtavascular microglia at P1-P5 (46.3% at P1 and 44.4% at P5) in the frontal cortex and at P5-P7 (39.1% at P5 and 34.2% at P7) in the more caudal somatosensory cortex (Fig. 3B-C, line graphs). Moreover, during times of active microglial colonization in both postnatal cortical regions (P1-P5 in the frontal cortex and P1-P7 in the somatosensory cortex), significantly more microglial primary processes were aligned parallel with vessels compared to older ages (Fig. 3D-G). This parallel juxtavascular microglial orientation along vessels is consistent with a migratory orientation.

**Figure 3:**
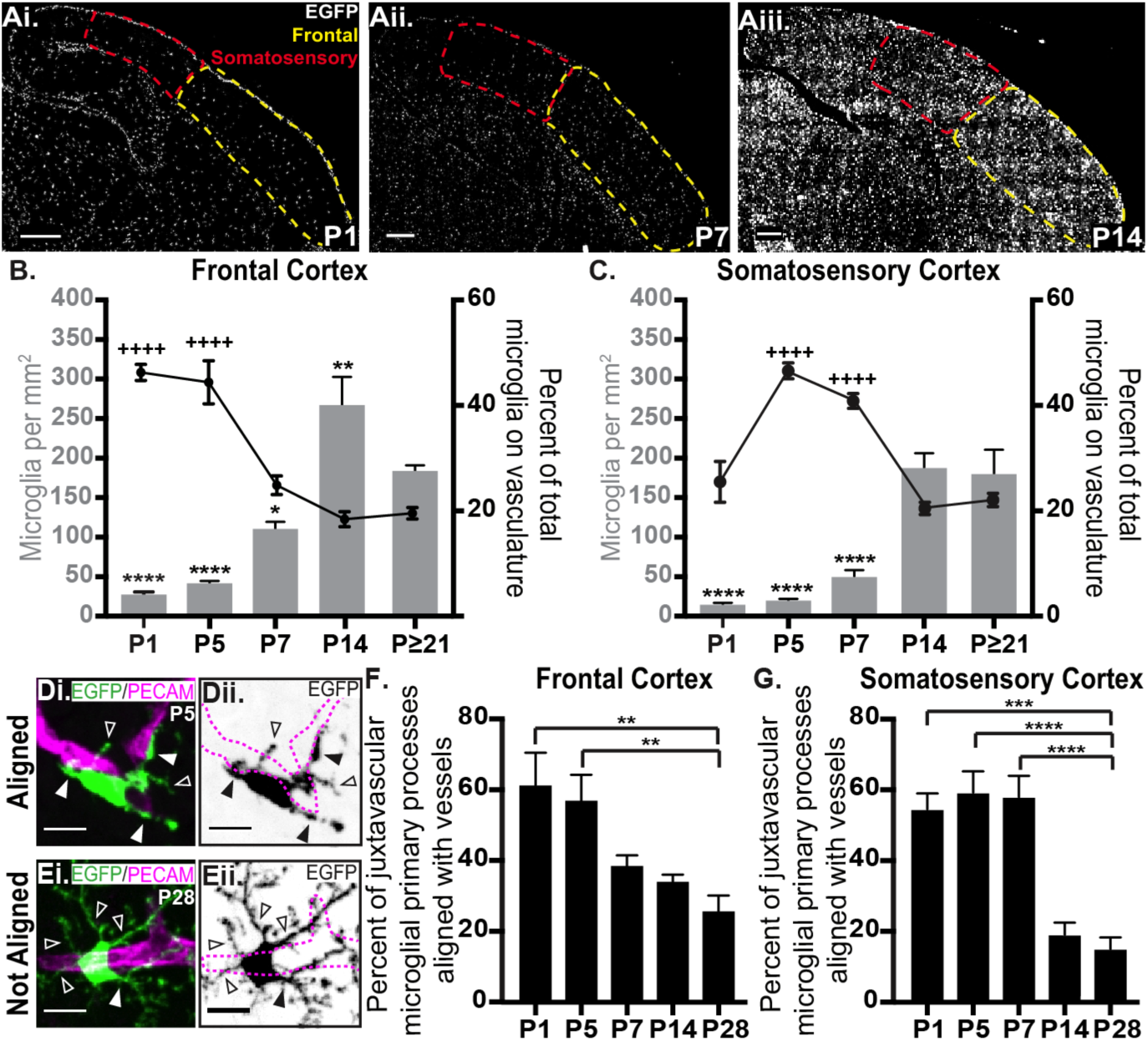
Microglia associate and align with vasculature as they colonize the cortex in a rostral-to-caudal gradient. **A.** Tiled sagittal sections of a P1 (**Ai**), P7 (**Aii**), and P14 (**Aiii.**) *Cx3cr1^EGFP/+^* brain. The dotted yellow and red lines outline the frontal and somatosensory cortex, respectively. Scale bars= 400µm. **B-C.** Left Y axis and gray bars: quantification of microglial density over development in the frontal cortex (**B**) and somatosensory cortex (**C**). One-way ANOVA with Dunnett’s post hoc; comparison to P≥21, n=4 littermates per developmental time point, *p<.05, **p<.01, ****p<.0001. Right Y axis and black line graphs: the percent of the total microglia population associated with vasculature over development in the frontal cortex (**B**) and somatosensory cortex (**C**). Note, data corresponding to the percent of juxtavascular microglia in the frontal cortex (line graph in C) are the same as presented in Fig. 1G. One-way ANOVA with Dunnett’s post hoc; comparison to P≥21, n=4 littermates per developmental time point, ++++p<.0001. **D-E.** Representative images of juxtavascular microglia (EGFP, green in **Di** and **Ei**; black in **Dii** and **Eii**) primary processes aligned parallel (**D**) with vessels (magenta, anti-PECAM) in the P5 frontal cortex, which were largely not aligned at P28 (**E**). Filled arrowheads denote processes aligned parallel to the vessel and unfilled arrowheads denote those microglial processes that are not aligned with the vessel. The dotted magenta line in **Dii** and **Eii** outline the vessel in **Di** and **Ei**. Scale bars= 10µm. **F-G.** Quantification of the percent of juxtavascular primary processes that are aligned parallel with vessels in the frontal (**F**) and somatosensory (**G**) cortices over development. One-way ANOVA with Dunnett’s post hoc; comparison to P≥21, n=3-4 littermates per developmental time point, *** p<.001, ****p<.0001. All error bars represent ± SEM.

To further investigate microglia-vascular interactions in the context of colonization of the postnatal cortex, we assessed a somatosensory sub-region where the pattern of microglial colonization has been well described—the barrel cortex. Layer IV of the barrel cortex contains thalamocortical synapses, which form a highly precise synaptic map of the vibrissae (whiskers) on the snout. These layer IV thalamocortical synapses form discrete barrel structures corresponding to each whisker, which are separated by septa where thalamocortical synapses are largely absent (Fig. 4A) (Woolsey and Van der Loos 1970; Welker and Woolsey 1974). Previous work has shown that microglia first localize to the septa and then colonize these thalamocortical synapse-dense barrel centers between P6 and P7 and this process is delayed to P8-P9 day in CX3CR1-deficient (*Cx3cr1^-/-^*) mice (Hoshiko et al. 2012). This delay in recruitment in *Cx3cr1^-/-^* mice is concomitant with a delay in synapse maturation. However, it was unclear how CX3CR1 was regulating the timing of microglial recruitment to synapses in the barrel cortex. To identify barrels, we labeled thalamocortical presynaptic terminals with an antibody against vesicular glutamate transporter 2 (VGluT2). Microglia were labeled with transgenic expression of EGFP in either *Cx3cr1^+/-^* (*Cx3cr1^EGFP/+^*) or *Cx3cr1^-/-^* (*Cx3cr1^EGFP/EGFP^*) mice. The vasculature was labeled with anti-PECAM. Similar to previous work (Hoshiko et al. 2012), microglia infiltrated thalamocortical synapse-dense barrel centers (outlined with a yellow dotted line in Fig. 4C-F) from the septa by P6-P7 in *Cx3cr1^+/-^* mice and this process was delayed by one day in *Cx3cr1^-/-^* mice (Fig. 4B-D). Strikingly, just prior to entering barrel centers at P5-P6 in *Cx3cr1^+/-^* mice, a higher percentage of microglia were juxtavascular (Fig. 4E, G, arrowheads). Further, this microglia-vascular association was delayed by one day in *Cx3cr1^-/-^* mice (Fig. 4F-G), which is consistent with the delay in microglial migration into barrel centers in these mice (Fig. 4B). In both genotypes, the percentage of juxtavascular microglia decreased once the microglia began to colonize the thalamocortical synapse-dense barrel centers, P7 in *Cx3cr1^+/-^* mice and P8 in *Cx3cr1^-/-^* mice (Fig. 4F-G). These changes in microglia-vascular interactions were independent of any changes in total microglial or vascular density in layer IV (Fig. 4H-I), but rather specific to microglial distribution between the septa and barrels. These data are consistent with a model by which microglia use the vasculature to colonize synapse-dense cortical regions at the appropriate developmental timing. They further suggest that CX3CR1 signaling modulates the timing of microglial-vascular interactions and, subsequently, colonization to synapse-dense regions of the barrel cortex.

**Figure 4:**
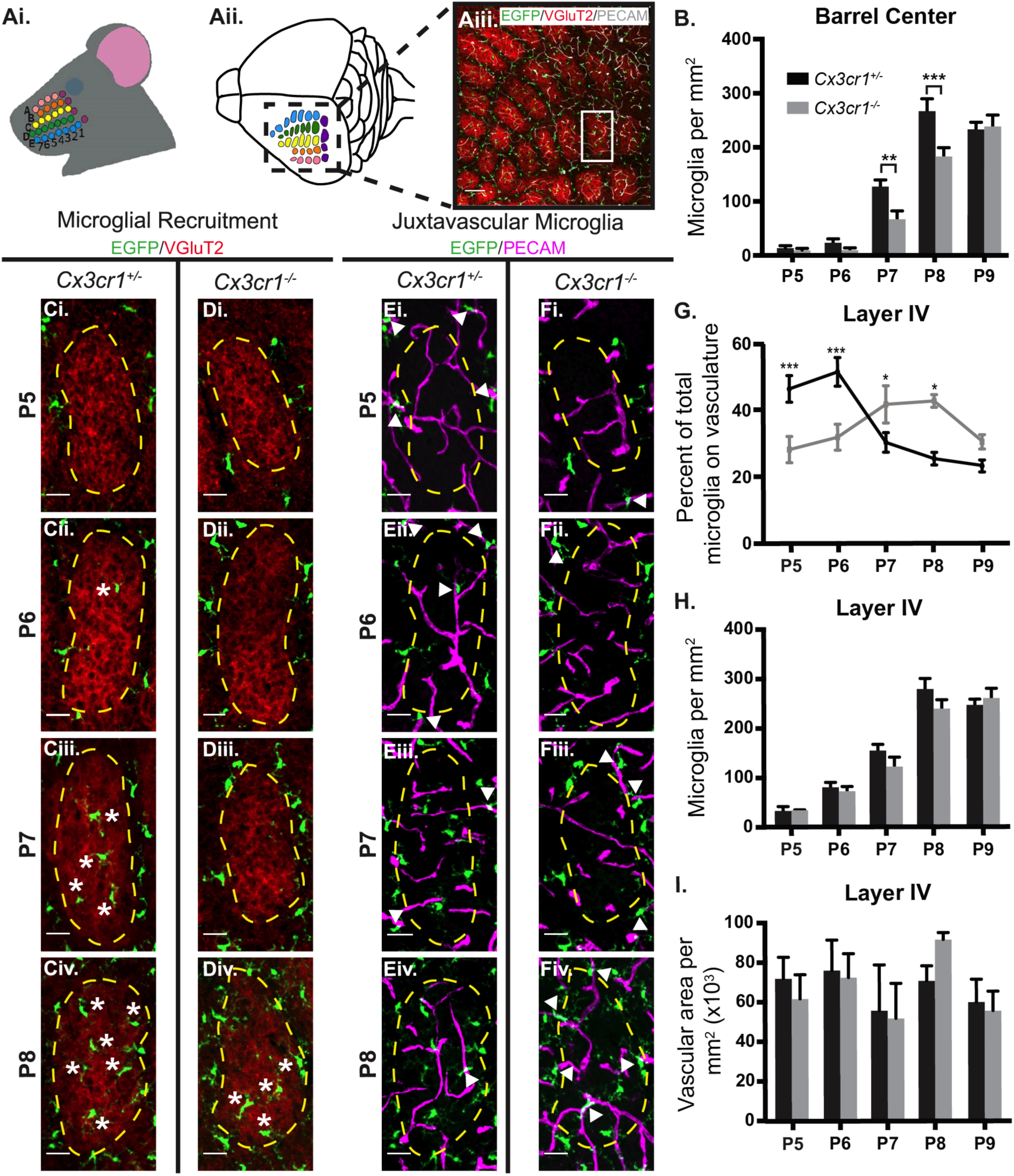
A high percentage of microglia associate with vasculature as they are recruited to synapses in the cortex in a CX3CR1-dependent manner. **Ai-Aii.** Layer IV of the barrel cortex contains thalamocortical synapses, which form a highly precise synaptic map of the vibrissae (whiskers) on the snout. **Aiii.** A low magnification representative image of a tangential section through layer IV of the barrel cortex shows layer IV thalamocortical presynaptic terminals (red, anti-VGluT2), form discrete barrel structures corresponding to each whisker, which are separated by septa where thalamocortical terminals are largely absent. Microglia are labeled by EGFP (green) and the vasculature is labeled with anti-PECAM (gray). White box denotes a single barrel. Scale bar= 100µm. **B**. Quantification of the number of microglia per mm^2^ within the barrel centers in developing *Cx3cr1^+/-^* (black bars) and *Cx3cr1^-/-^* (gray bars) mice. Two-way ANOVA with a Sidak’s post hoc; n=4 littermates per genotype per developmental time point; ** p<.01, ***p<.001. **C-D.** Representative images of quantification in B. Images are zoomed in to show single barrels within tangential sections of layer IV of the barrel cortex (denoted by white box in Aiii) where microglia (green) are recruited to barrel centers in *Cx3cr1^+/-^* by P7 (**C**) and in *Cx3cr1^-/-^* by P8 (**D**). Asterisks denote microglia located within barrel centers. The dotted yellow lines denote the perimeters of the VGluT2-positive thalamocortical inputs (red), which define the barrels vs. the septa. Scale bars= 30µm. **E-F**. The same representative fields of view in C-D but lacking the anti-VGluT2 channel and, instead, including the channel with anti-PECAM immunostaining (magenta) to label the vessels. Microglia are still labeled with EGFP (green). Dotted yellow lines still denote the perimeters ofthe VGluT2-posiive barrels (red in C-D). Juxtavascular microglia in *Cx3cr1^+/-^* and *Cx3cr1^-/-^* mice are denoted by filled arrowheads. Scale bar= 30µm. **G.** Quantification of the percent of microglia associated with the vasculature in *Cx3cr1^+/-^* (black lines) and *Cx3cr1^-/–^* (gray lines) animals over development in layer IV of the barrel cortex demonstrates a peak of vascular association in *Cx3cr1^+/-^* mice at P5-P6, which is delayed to P7-P8 in *Cx3cr1^-/-^* coincident with delayed microglial recruitment to barrel centers. Two-way ANOVA with a Tukey’s post hoc; n=4-5 littermates per genotype per developmental time point; *p<.05, ***p<.001, compared to P9 *Cx3cr1^+/-^*. **H-I.** Quantification of microglial (**H**) and vascular (**I**) density in *Cx3cr1^+/-^* (black bars) and *Cx3cr1^-/-^* (gray bars) animals over development in layer IV of the barrel cortex. Two-way ANOVA with a Sidak’s post hoc; n=4 littermates per genotype per developmental time point. All error bars represent ± SEM.

### Juxtavascular microglia migrate along the vasculature as they colonize the developing brain and are stationary in adulthood

With data demonstrating that high percentages of microglia are juxtavascular when they are actively colonizing the brain with processes aligned parallel to the vessel, we next performed live imaging to assess migration. As the early postnatal cortex is challenging to image *in vivo*, we performed our initial analyses in acute cortical slices. Acute slices of somatosensory cortex were prepared from early postnatal (P7) and adult (P≥120) *Cx3cr1^EGFP/+^* mice, which were given a retro-orbital injection of Texas Red labeled dextran to label blood vessels prior to slice preparation. We then imaged microglia every 5 minutes over 6 hours at both ages (Fig. 5A). Live imaging at P7 revealed significant juxtavascular microglial soma movement along blood vessels in the somatosensory cortex compared to vascular-unassociated microglia at P7 (Fig. 5B, D, see also Movies 3-5). Specifically, 28.6% of juxtavascular microglia somas moved at a rate of 3-5µm/hour and another 26.1% moved at a rate of 5-7.5µm/hour (Fig. 5D). In comparison, only 9.3% and 6.8% of vascular-unassociated microglia at the same age moved at 3-5µm/hour and 5-7.5µm/hour, respectively. We further found that when we assessed just the motile soma at P7, significantly more juxtavascular microglia somas travelled >20µm (30.9% traveled 20-30µm and 23.6% traveled 30-45µm) over 6 hours compared to vascular-unassociated microglia (7.5% and 6.8% traveled 20-30µm and 30-45µm, respectively) (Fig. 5E). Importantly, the juxtavascular microglia soma velocities and distances traveled are consistent with the rate and distances at which microglia migrate to barrel centers within the somatosensory cortex *in vivo* where the distance between the septa and barrel center is ∼80µm and it takes ∼24 hours for microglia to reach the barrel center from the septa. Demonstrating directional motility and suggesting migration along the vessel, 84.1% of these postnatal juxtavascular microglia had a motility trajectory of ≤15° along the blood vessel (Fig. 5F). Together, these data demonstrate directional migration of juxtavascular microglia at distances and speeds consistent with colonization of the cortex (P7).

**Figure 5:**
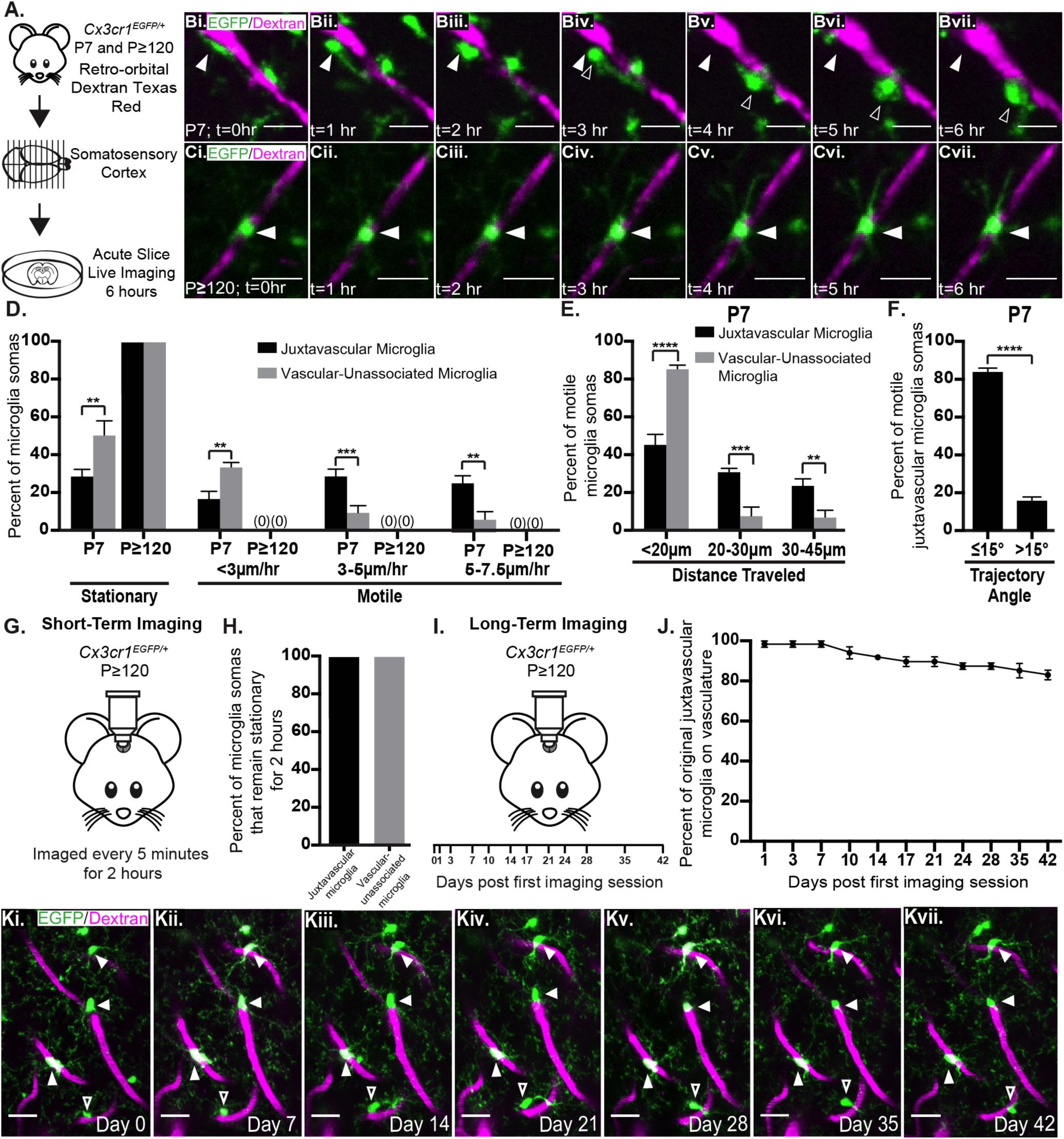
Juxtavascular microglia migrate along blood vessels as they colonize the developing brain and are largely stationary in adulthood. **A.** A schematic of the live imaging experiment. *Cx3cr1^EGFP/+^* mice received a retro-orbital injection of Texas red labeled dextran to label the vasculature 10 minutes prior to euthanasia. Coronal somatosensory cortices were cut and imaged every 5 minutes over 6 hours immediately following slice preparation. **B-C**. Representative fluorescent images from a 6-hour live imaging session from a P7 (**B**) and P≥120 (**C**) slice. Filled arrowheads indicate microglial soma position at t=0. Unfilled arrowheads indicate the location of the same microglial soma at 0hr (**Bi, Ci**), 1hrs (**Bii, Cii**), 2hrs (**Biii, Ciii**), 3hrs (**Biv, Civ**), 4hrs (**Bv, Cv**), 5hrs (**Bvi, Cvi**), and 6hrs (**Bvii, Cvii**). See also Movies 3-6. Scale bars= 30µm. **D**. Quantification of juxtavascular (black bars) and vascular-unassociated (gray bars) microglia soma motility speed/velocity. Two-way ANOVA with a Sidak’s post hoc; n=4 mice per time point; **p<.01, ***p<.001. **E.** Quantification of the distance traveled of juxtavascular (black bars) and vascular-unassociated (gray bars) microglia somas in the P7 somatosensory cortex. Two-way ANOVA with a Sidak’s post hoc; n=4 mice; **p<.01, ***p<.001, ****p<.0001. **F.** Quantification of migratory juxtavascular microglia trajectory angles in the P7 somatosensory cortex. Unpaired student’s t-test; n=4 mice per time point; ****p<.0001. **G.** A schematic of short-term 2-photon live imaging experiment in adult cortex. *Cx3cr1^EGFP/+^* mice received a retro-orbital injection of Texas Red-labeled dextran to visualize the vasculature 10 min prior to each imaging session. EGFP+ juxtavascular microglia were then imaged every 5 minutes for 2 hours. See also Movie 7. **H.** Quantification of the percent of juxtavascular (black bars) and vascular-unassociated (gray bars) microglia that remain stationary for 2 hours. Unpaired student’s t-test; n=3 mice per developmental time point. **I.** A schematic of the long-term 2-photon live imaging experiment in adult visual cortex. *Cx3cr1^EGFP/+^* mice received a retro-orbital injection of Texas Red-labeled dextran to visualize the vasculature 10 min prior to each imaging session. EGFP+ juxtavascular microglia were then imaged for 6 weeks. **J.** Quantification of the percent of juxtavascular microglia on vessels on day 0 that remain on vessels through six weeks of imaging. Data are representative of n=3 mice. **K.** Representative fluorescent images acquired during a 6-week live imaging session from a single mouse. Filled arrowheads indicate juxtavascular microglia that remain on vessels for 6 weeks. Unfilled arrowhead indicates a juxtavascular microglia that changes position, but remains on the vasculature, over 6 weeks. All error bars represent ± SEM.

Interestingly, this migratory behavior along the vasculature was developmentally regulated and juxtavascular microglia in adult slices were largely stationary (Fig. 5C-D; see also Movie 6). We further confirmed the stationary phenotype of juxtavascular microglia in the adult cortex by *in-vivo* 2-photon live imaging in *Cx3cr1^EGFP/+^* mice. Windows were placed over the visual cortex, which was most conducive to our head posts necessary for stabilizing the head in awake, behaving mice during imaging. We have found similar microglia-vascular interactions by static confocal imaging in the visual cortex (data not shown). Mice were given a retro-orbital injection of Texas Red labeled dextran to label blood vessels prior to imaging and juxtavascular microglia were imaged every 5 min over the course of 2 hours (Fig. 5G). As observed in acute cortical slices, 100% of juxtavascular and vascular-unassociated microglia were stationary (Fig. 5H; see also Movie 7). To further understand long-term dynamics, we imaged juxtavascular microglia *in vivo* over the course of 6 weeks (Fig. 5I). We identified that 82.9% of juxtavascular microglia present on day 0 of imaging remained on the vasculature 6 weeks later (Fig. 5J-K). Together, these data demonstrate that juxtavascular microglia in the postnatal cortex are highly migratory compared to non-vascular associated microglia. In contrast, juxtavascular microglia in adulthood are largely stationary, which suggests the establishment of a niche for juxtavascular microglia in the adult brain.

### Microglia associate with the vasculature in areas lacking full astrocyte endfoot coverage

Our data demonstrate a strong microglial association and migration along the developing postnatal cortical vasculature. One possible mechanism regulating these developmental changes in juxtavascular microglia is the changing cellular composition of the NVU over development. The neurovascular unit (NVU) begins to form during embryonic development, when pericytes associate with endothelial cells. Later in postnatal development, astrocytes are born and begin wrapping their endfeet around vessels until the vast majority of the vasculature is ensheathed by astrocyte endfeet by adulthood (Daneman et al. 2010; Schiweck, Eickholt, and Murk 2018; Bayraktar et al. 2015). As previously described (Daneman et al. 2010), the territory of Aquaporin 4 (AQP4)-positive astrocyte endfeet on PDGFRβ+ capillaries was low in the early postnatal cortex and then expanded over the first postnatal week (Fig. 6 A-D, bar graph in D). In more mature animals (≥P21), astrocytic endfeet covered ∼85% of vessels in the frontal cortex. Intriguingly, this developmental timing of astrocyte endfoot coverage mirrored the developmental shift in the percentage of juxtavascular microglia in the cortex (Fig. 6D, line graph). That is, as the percentage of juxtavascular microglia decreased, astrocyte endfoot coverage increased. This astrocyte coverage also correlated with the timing of decreased microglial motility along the vessels (Fig. 5). We further assessed microglia-astrocyte endfoot interactions by confocal microscopy and 3D surface rendering. At all ages, microglia only contacted the vasculature in areas either completely void of astrocyte endfeet or in areas where vessels were not fully covered by the endfeet (Fig. 6 A-C, white arrow heads; Fig. 6E, see also Movies 8-10). Juxtavascular microglia were never in direct contact with solely astrocyte endfeet and no vessel at any age assessed (Fig. 6E). Given that cells of the NVU are nanometers apart from each other, we confirmed these results with expansion microscopy (ExM; Fig 6F-G), structured illumination microscopy (SIM; Fig. 6H-I) and electron microscopy (EM; Fig 7). By EM, microglia were identified based on characteristic microglial morphologies. Microglia nuclei tend to be half-mooned shape or long and thin with electron dense heterochromatin around the edge of the nucleus. Microglia were further distinguished by EM from perivascular macrophages by having processes emanating from the soma. Serial sectioning and 3D reconstruction of a representative cell captured by EM from each age confirmed that juxtavascular microglia contacted the basal lamina in vascular areas without full astrocyte endfoot coverage at all ages (Fig. 7C, see also Movies 11 and 12). Together, these data demonstrate that juxtavascular microglia associate with the vascular basal lamina and associate with the vasculature in areas lacking full coverage by astrocyte endfeet. The data raise the intriguing possibility that lack of astrocyte endfeet in early postnatal development provides a permissive environment for juxtavascular microglial association with and migration along the vasculature as they colonize the brain.

**Figure 6:**
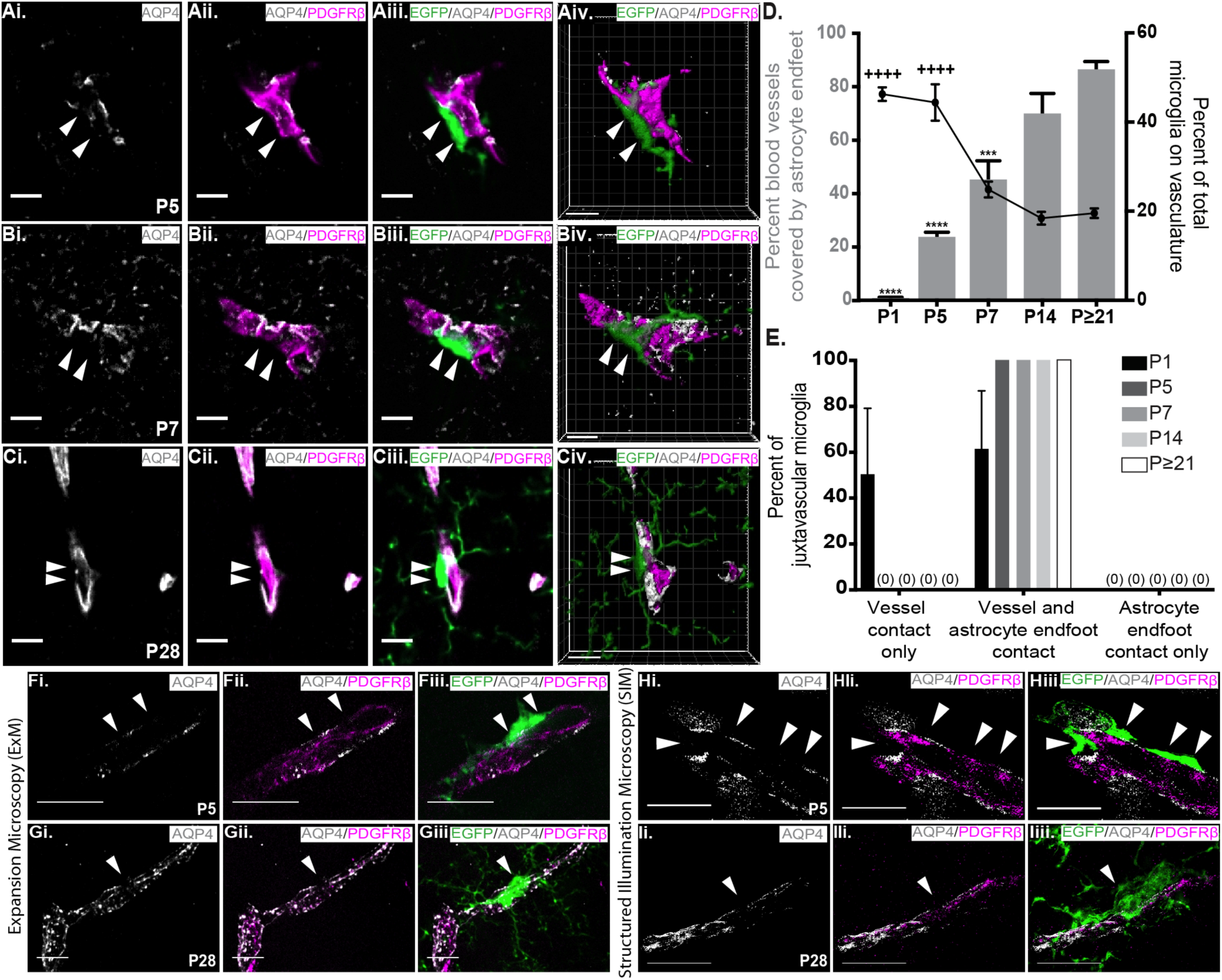
Juxtavascular microglia contact the cortical vasculature in areas lacking full astrocytic endfoot coverage. **A-C.** Representative single optical plane images and 3D rendering (**Aiv-Civ**; see also Movies 8-10) of juxtavascular microglia (green, EGFP) and blood vessels (magenta, anti-PDGFRβ) in areas void of astrocytic endfoot labeling (gray, anti-AQP4) in the frontal cortex at P5 (**A**), P7 (**B**) P28 (**C**). Filled arrowheads denote vascular areas that lack astrocyte endfeet where juxtavascular microglia are contacting the vessel. Scale bars= 10µm. **D.** Left Y axis, gray bars: quantification of the percent of blood vessels covered by astrocyte endfeet over development in the frontal cortex. One-way ANOVA with Dunnett’s post hoc; comparison to P≥21, n=3 littermates per developmental time point, ***p<.001, ****p<.0001. Right Y axis, black line: the percent of the total microglia population that are juxtavascular over development in the frontal cortex (data are the same as presented in Fig. 1G). One-way ANOVA with Dunnett’s post hoc; comparison to P≥21, n=4 littermates per developmental time point, ++++p<.0001. **E.** Quantification of the percent of juxtavascular microglia contacting vessels only, vessels and astrocyte endfeet (representative images in A-C), and astrocyte endfeet only from 3D rendered images. **F-I.** Representative expansion microscopy (ExM, **F-G**) and structured illumination microscopy (SIM, **H-I**) images of juxtavascular microglia (green, EGFP), in vascular areas lacking anti-AQP4 (gray) astrocytic endfoot labeling (filled arrowheads) in the P5 (**F, H**) and P28 (**G, I**) frontal cortex. Scale bars= 10µm. All error bars represent ± SEM.

**Figure 7:**
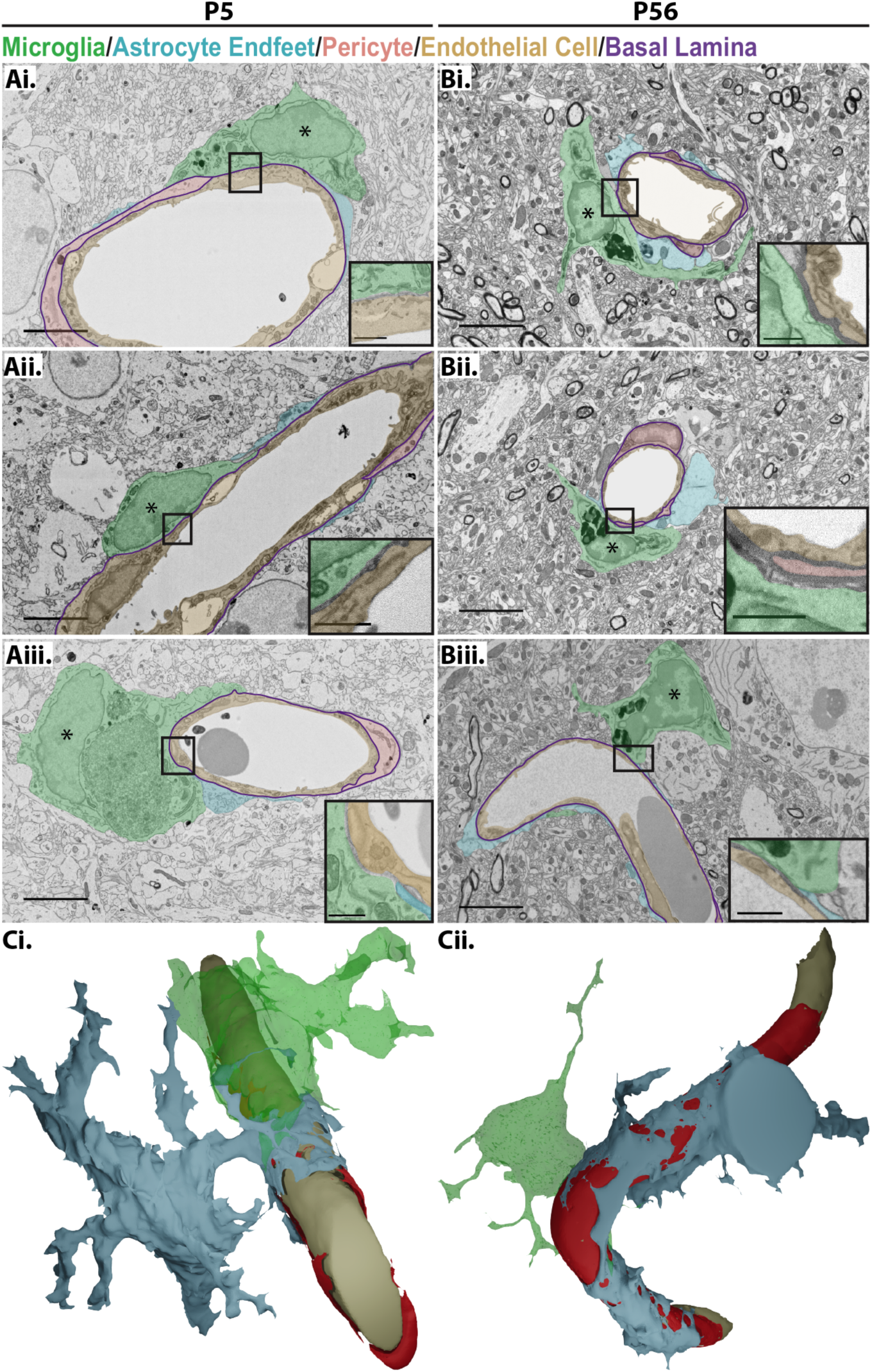
Ultrastructural analysis by EM reveals that juxtavascular microglia directly contact the basal lamina of the vasculature. **A-B.** Electron microscopy (EM) of juxtavascular microglia (green pseudocoloring) contacting the basal lamina (purple line) of a blood vessel in an area void of astrocyte endfeet (blue pseudocoloring) in the P5 (**A**, left column) and P56 (**B**, right column) frontal cortex. Pink pseudocoloring denotes a pericyte. Asterisks denote microglia nuclei. Scale bar= 5µm. The black box denotes the magnified inset in the bottom right corner where microglia (green pseudocoloring) directly contact the basal lamina (unlabeled in the inset) and only partially contacts the astrocyte endfoot (blue pseudocoloring). Scale bar= 1µm. **C.** 3D reconstruction of serial EM of P5 juxtavascular microglia in **Aiii** (**Ci**) and P56 P56 juxtavascular microglia in **Biii** (**Cii**) (see also Movies 11 and 12).

## DISCUSSION

This study provides the first extensive analysis of juxtavascular microglia in the healthy developing and adult brain. We discovered that a high percentage of juxtavascular microglia are in direct contact with largely capillaries in the early postnatal mouse cortex. Similar microglia-vascular association was observed in the developing human brain. Live imaging revealed that juxtavascular microglia are migratory along the vasculature during the peak of microglial colonization of the postnatal cortex and become stationary by adulthood. In addition, microglia are highly associated with the vasculature during development as they are being recruited to synapse-dense rich cortical regions and the timing of these interactions is regulated by CX3CR1. Last, we provide evidence that microglia preferentially contact the vasculature at all ages in areas lacking full astrocyte endfoot coverage and expansion of astrocytic endfeet along blood vessels coincides with a decrease in microglia migration along vessels. Taken together, these data suggest that microglia are using the vasculature to migrate and colonize the cortex and the timing of this vascular association is CX3CR1-dependent. Our data further support a mechanism in which microglial migration along the vasculature during development ceases and juxtavascular microglia become stationary upon the maturation of astrocyte endfeet.

### A possible role for the vasculature in regulating microglial colonization

Microglia are born as primitive macrophages in the embryonic yolk sac and enter the neuroepithelium at embryonic day E9.5 by crossing the pial surface and lateral ventricles (Navascués et al. 2000; Swinnen et al. 2013; Ginhoux et al. 2010). Microglia then migrate and proliferate through the brain parenchyma in a rostral-to-caudal gradient to colonize the embryonic brain (Sorokin et al. 1992; Navascués et al. 2000; Swinnen et al. 2013; Alliot, Godin, and Pessac 1999; Perry, Hume, and Gordon 1985; Ashwell 1991). Signaling mechanisms have been identified to regulate initial microglial infiltration into the brain parenchyma, such as matrix metalloproteinases (MMPs), stromal cell derived factor 1 (SDF-1), and Cxcl12/Cxcr4 signaling (Ginhoux et al. 2010; Arno et al. 2014; Ueno and Yamashita 2014). However, far less is known about the mechanisms regulating microglial localization to the appropriate brain regions once they reach the parenchyma, particularly during postnatal development. Previous work has shown microglia can migrate along the vasculature in acute embryonic brain slices and brain slices prepared from postnatal mice in an injury context (Smolders et al. 2017; Grossmann et al. 2002). In addition, other work has shown that oligodendrocyte precursor cells (OPCs) require the vasculature as a physical substrate for migration (Tsai et al. 2016). Similar findings have been identified for neural stem cells where the timing of astrocyte endfeet to the vessels has also been implicated (Bovetti et al. 2007; Fujioka, Kaneko, and Sawamoto 2019; Whitman et al. 2009). We have identified that microglia are highly associated with vasculature during the peak of microglial colonization and recruitment to synapses. Furthermore, these vascular-associated microglia are migratory along blood vessels during early postnatal development and later become stationary once microglial colonization is complete. We also show in CX3CR1-deficient mice with known delays in microglial colonization of synapse-dense cortical regions that there are concomitant delays in microglial association with the vasculature. As we have observed no significant expression of *Cx3cl1* (the CX3CR1 ligand) by vascular cells (Gunner et al. 2019) and a subset of microglia still associate with the vasculature in *Cx3cr1^-/-^* mice, this delay in microglial vascular association in *Cx3cr1^-/-^* mice is most likely due to disruptions in chemokine gradient signaling from neuronal sources of CX3CL1 versus a direct effect of vascular adhesion. This would suggest that microglia receive directional cues from surrounding cells, use the vasculature as a physical substrate to migrate towards those cues, and the timing of this migration along the vasculature is regulated by CX3CR1. As *Cx3cr1^-/-^* mice have delays in synapse maturation and pruning and, long-term, have behavioral deficits consistent with an autism-like phenotype, it suggests that these microglia-vascular associations in development have long-term consequences (Paolicelli et al. 2011; Zhan et al. 2014; Hoshiko et al. 2012). The vascular cues regulating microglial adhesion and migration in the healthy CNS are yet to be identified, which will be key to determine the relative importance of microglia-vascular interactions for microglial colonization, brain development, and long-term CNS function.

### Microglia-astrocyte interactions at the NVU interface

Another interesting direction is to determine the role of astrocyte endfeet in regulating microglia-vascular interactions. Astrocytes are born and begin wrapping their processes to form endfeet along blood vessels during the first postnatal week (Daneman et al. 2010). By adulthood, astrocytes endfeet ensheath 60-95% of the vasculature (Mathiisen et al. 2010; Korogod, Petersen, and Knott 2015). Here, we demonstrate that juxtavascular microglia in the postnatal cortex represent a large percentage of total microglia and are migratory along the vasculature. Juxtavascular microglia migration decreases as astrocyte endfeet develop and ensheath the vasculature. In addition, we showed that microglia contact vessels at all ages in areas lacking full astrocytic endfoot coverage and EM revealed high association between juxtavascular microglia and the vascular basal lamina. These data raise the intriguing possibility that the basal lamina provides an adhesive substrate for microglial association and migration, which becomes restricted upon astrocyte endfoot arrival. Astrocyte endfeet may, therefore, physically exclude microglia from contacting the basal lamina and associating with the vasculature. Another possibility is that microglia in the postnatal brain repel astrocyte endfeet, but this repellent signal later decreases as the animal matures so that astrocyte endfeet can wrap the vessels. Analysis of astrocyte endfoot-juxtavascular microglia interactions along blood vessels will be important going forward.

### Possible functions for juxtavascular microglia in the healthy CNS

Are juxtavascular microglia a unique subpopulation of microglial cells that perform distinct functions at the NVU? Evidence in the literature suggests microglia play important roles in regulating the vasculature, but it is unclear if these functions are specific to juxtavascular microglia. For example, in the embryonic brain, microglia are often localized to vascular junction points and depletion of all microglia is associated with a decrease in vascular complexity (Fantin et al. 2010). Similar findings have been identified in the developing retina (Rymo et al. 2011; Checchin et al. 2006; Dudiki et al. 2020). Our data demonstrating that microglia are localized to the vasculature prior to the arrival of the astrocyte endfeet could place microglia in a position to regulate fine-scale remodeling of the vasculature throughout the brain and/or help to maintain the BBB prior to astrocyte endfoot arrival. Arguing against the latter, microglia depletion during development does not appear to induce changes in BBB integrity in the postnatal brain (Parkhurst et al. 2013; Elmore et al. 2014). These data are in contrast to the inflamed adult CNS, were microglia regulate BBB integrity (Zhao et al. 2018; Stankovic, Teodorczyk, and Ploen 2016). One of the most recent studies shows that during systemic inflammation, parenchymal microglia migrate to the vasculature and help to maintain the BBB at acute stages (Haruwaka et al. 2019). However, with sustained inflammation, microglia phagocytose astrocyte endfeet and facilitate BBB breakdown. In the absence of inflammation, it remains unknown what functions juxtavascular microglia may perform. However, our *in vivo* live imaging data demonstrating microglia in the adult brain are stationary for nearly 2 months opens up the possibility that these cells reside in a vascular niche to perform specialized functions. One possible role could be to serve as immune surveillant “first responders”. We report that microglia are in direct contact with the basal lamina at all ages in areas lacking astrocyte endfeet, which typically work to maintain the BBB (Abbott, Rönnbäck, and Hansson 2006; Kimelberg and Nedergaard 2010; Macvicar and Newman 2015). As a result, juxtavascular microglia in the healthy brain are in an ideal location to serve as conduits to relay changes in the peripheral environment (changes in the microbiome, infection, etc.) to the CNS. Indeed, many of these changes in peripheral immunity are known to directly impact neural circuit function and behavior in ways we do not yet fully understand (Hanamsagar and Bilbo 2017; Lebovitz et al. 2018; Abdel-haq et al. 2018) and recent work in the inflamed brain has demonstrated microglia can serve as a conduit by which the microbiome affects neuroinflammation (Rothhammer et al. 2018).

### Microglia-vascular interactions: Implications for CNS disease

Our findings have important implications for neurological diseases associated with the injured or aged CNS where there is enhanced microglial association with the vasculature, such as in stroke, brain tumors, multiple sclerosis (MS), and Alzheimer’s disease (Stankovic, Teodorczyk, and Ploen 2016; Zhao et al. 2018). This enhanced association can lead to further breakdown of the BBB and infiltration of peripheral immune cells into the CNS and possibly angiogenesis in the case of brain tumors (Stankovic, Teodorczyk, and Ploen 2016; Zhao et al. 2018; Haruwaka et al. 2019). Therefore, understanding precisely when and where microglia interact with the vasculature in the healthy brain may lead to therapeutic strategies to reduce vascular pathology and facilitate recovery. One intriguing possibility is that these sites of microglial contact, which lack astrocyte endfeet, are more vulnerable to BBB breakdown and infiltration of peripheral immune cells and factors. In addition to neurodegenerative disorders, our findings may also have important implications for neurodevelopmental disorders such as autism spectrum disorders (ASDs). For example, microglia-vascular interactions may be important for the timing of microglial colonization to synapse-dense brain regions where they regulate synapse maturation and pruning during critical windows in development (Paolicelli et al. 2011; Hoshiko et al. 2012; Tremblay, Lowery, and Majewska 2010; Schafer et al. 2012; Gunner et al. 2019). If these interactions are disrupted, the timing of synapse development and, ultimately, neural circuit function may be altered. This is supported by our data from *Cx3cr1^-/-^* mice showing delays in microglial association with the vessels, which is concomitant with known delays in microglial recruitment to developing synapses and delays in synapse maturation in these mice (Paolicelli et al. 2011; Zhan et al. 2014; Hoshiko et al. 2012). Long term, *Cx3cr1^-/-^* mice have phenotypes associated with ASD, including decreased functional brain connectivity, deficits in social interactions, and increased repetitive behaviors (Zhan et al. 2014). However, a better understanding of how vascular interactions affect microglial colonization and extending these analyses of microglia-vascular interactions into the ASD human brain will be necessary.

Together, our work sheds new light on an understudied population of microglia, juxtavascular microglia. This work lays the foundation for identifying new molecular mechanisms underlying microglia-vascular interactions, identifying mechanistic underpinnings of microglia-astrocyte crosstalk at the level of the NVU, and furthering our understanding of juxtavascular microglia function in CNS homeostasis. With the vascular interface emerging as an important aspect of many neurological conditions, this study also lays the critical groundwork to study how this microglial population may be important in a wide range of CNS diseases.

## Acknowledgements

We thank Oleg Butovsky (Brigham and Women’s Hospital, Harvard University) for providing the anti-P2RY12 antibody and Georg Kislinger for help with electron microscopy. This work was funded by NIMH-R01MH113743 (DPS), NINDS-P01NS083513 (EJH), NIAID-T32 A1095213 (EM), AHA Predoctoral Fellowship 19PRE3480616 (JC), Brain & Behavior Research Foundation (DPS), Charles H. Hood Foundation (DPS), the Dr. Miriam and Sheldon G. Adelson Medical Research Foundation (DPS and MS), Deutsche Forschungsgemeinschaft (DFG, German Research Foundation) under Germany’s Excellence Strategy within the framework of the Munich Cluster for Systems Neurology (EXC 2145 SyNergy – ID 390857198, MS).

---

**Movie 1: 3D rendering of juxtavascular microglia in the early postnatal frontal cortex.** 3D reconstruction and surface rendering of juxtavascular microglia (green, EGFP) associated with blood vessels (magenta, anti-PECAM) in the P5 frontal cortex. Yellow denotes contact area between microglia and blood vessels.

**Movie 2: 3D rendering of juxtavascular microglia in the P28 frontal cortex.** 3D reconstruction and surface rendering of juxtavascular microglia (green, EGFP) associated with blood vessels (magenta, anti-PECAM) in the P28 frontal cortex. Yellow denotes contact area between microglia and blood vessels.

**Movie 3: Juxtavascular microglial migration in the early postnatal somatosensory cortex.** Representative live imaging of juxtavascular microglia (green, EGFP) migrating on vessels (magenta; dextran) in the P7 somatosensory. *Cx3cr1^EGFP/+^* mice received a retro-orbital injection of Texas red labeled dextran to label the vasculature 10 minutes prior to euthanasia. Coronal somatosensory cortices were imaged every 5 minutes over 6 hours immediately following slice preparation. Asterisk in still image denotes the microglia that was tracked for quantification.

**Movie 4: Juxtavascular microglial migration in the early postnatal somatosensory cortex.** A second representative live imaging of juxtavascular microglia (green, EGFP) migrating on vessels (magenta; dextran) in the P7 somatosensory. *Cx3cr1^EGFP/+^* mice received a retro-orbital injection of Texas red labeled dextran to label the vasculature 10 minutes prior to euthanasia. Coronal somatosensory cortices were imaged every 5 minutes over 6 hours immediately following slice preparation. Asterisk in still image denotes the microglia that was tracked for quantification.

**Movie 5: Juxtavascular microglial migration in the early postnatal somatosensory cortex.** A third representative live imaging of juxtavascular microglia (green, EGFP) migrating on vessels (magenta; dextran) in the P7 somatosensory. *Cx3cr1^EGFP/+^* mice received a retro-orbital injection of Texas red labeled dextran to label the vasculature 10 minutes prior to euthanasia. Coronal somatosensory cortices were imaged every 5 minutes over 6 hours immediately following slice preparation. Asterisk in still image denotes the microglia that was tracked for quantification.

**Movie 6: Juxtavascular microglial migration in the adult somatosensory cortex.** Representative live imaging of juxtavascular microglia (green, EGFP) stationary on vessels (magenta; dextran) in the P≥120 somatosensory cortex. *Cx3cr1^EGFP/+^* mice received a retro-orbital injection of Texas red labeled dextran to label the vasculature 10 minutes prior to euthanasia. Coronal somatosensory cortices were imaged every 5 minutes over 6 hours immediately following slice preparation. Asterisk in still image denotes the microglia that was tracked for quantification.

**Movie 7: 2-photon *in vivo* live imaging of juxtavascular microglia in the adult cortex.** Representative 2-photon *in* vivo live imaging of juxtavascular microglia (green, EGFP) stationary on blood vessels (magenta, dextran) over 2 hours in vivo in the adult cortex. *Cx3cr1^EGFP/+^* mice received a retro-orbital injection of Texas Red-labeled dextran to visualize the vasculature 10 min prior to each imaging session. EGFP+ juxtavascular microglia were then imaged every 5 minutes for 2 hours.

**Movie 8: Juxtavascular microglia contact the cortical vasculature in areas lacking full astrocytic endfoot coverage in the P5 frontal cortex.** 3D reconstruction and surface rendering of juxtavascular microglia (green, EGFP) contacting blood vessels (magenta, anti-PDGFRβ) in areas void of astrocytic endfoot labeling (gray, anti-AQP4) in the frontal cortex at P5.

**Movie 9: Juxtavascular microglia contact the cortical vasculature in areas lacking full astrocytic endfoot coverage in the P7 frontal cortex**. 3D reconstruction and surface rendering of juxtavascular microglia (green, EGFP) contacting blood vessels (magenta, anti-PDGFRβ) in areas void of astrocytic endfoot labeling (gray, anti-AQP4) in the frontal cortex at P7.

**Movie 10: Juxtavascular microglia contact the cortical vasculature in areas lacking full astrocytic endfoot coverage in the P28 frontal cortex**. 3D reconstruction and surface rendering of juxtavascular microglia (green, EGFP) contacting blood vessels (magenta, anti-PDGFRβ) in areas void of astrocytic endfoot labeling (gray, anti-AQP4) in the frontal cortex at P28.

**Movie 11: Serial EM 3D reconstruction of juxtavascular microglia in the early postnatal cortex.** 3D reconstruction of serial electron microscopy (EM) of juxtavascular microglia (green) contacting the basal lamina (red) of a blood vessel in an area void of astrocyte endfeet (blue) in the P5 frontal cortex.

**Movie 12: Serial EM 3D reconstruction of juxtavascular microglia in the P56 cortex.** 3D reconstruction of serial electron microscopy (EM) of juxtavascular microglia (green) contacting the basal lamina (red) of a blood vessel lacking full astrocyte endfoot (blue) coverage in the P56 frontal cortex.

## REFERENCES

Abbott, N. Joan, Lars Rönnbäck, and Elisabeth Hansson. 2006. “Astrocyte–Endothelial Interactions at the Blood–Brain Barrier.” Nature Reviews Neuroscience 7 (1): 41–53.

Abdel-haq, Reem, Johannes C M Schlachetzki, Christopher K Glass, and Sarkis K Mazmanian. 2018. “Microbiome – Microglia Connections via the Gut – Brain Axis.” Journal of Experimental Medicine 216 (1): 41–59.

Adams, Ryan A, Jan Bauer, Matthew J Flick, Shoana L Sikorski, Tal Nuriel, Hans Lassmann, Jay L Degen, and Katerina Akassoglou. 2007. “The Fibrin-Derived γ 377-395 Peptide Inhibits Microglia Activation and Suppresses Relapsing Paralysis in Central Nervous System Autoimmune Disease.” Journal of Experimental Medicine 204 (3): 571–82.

Alliot, Francoise, Isabelle Godin, and Bernard Pessac. 1999. “Microglia Derive from Progenitors, Originating from the Yolk Sac, and Which Proliferate in the Brain.” Developmental Brain Research 117: 145–52.

Armulik, Annika, Guillem Genové, Maarja Mäe, Maya H. Nisancioglu, Elisabet Wallgard, Colin Niaudet, Liqun He, et al. 2010. “Pericytes Regulate the Blood–Brain Barrier.” Nature 468 (7323): 557–61.

Arno, Benedetta, Francesca Grassivaro, Chiara Rossi, Andrea Bergamaschi, Valentina Castiglioni, Roberto Furlan, Melanie Greter, et al. 2014. “Neural Progenitor Cells Orchestrate Microglia Migration and Positioning into the Developing Cortex.” Nature Communications 5 (5611): 1–13.

Asano, Shoh M, Ruixuan Gao, Asmamaw T Wassie, Paul W Tillberg, Fei Chen, and Edward S Boyden. 2018. “Expansion Microscopy : Protocols for Imaging Proteins and RNA in Cells and Tissues.” Current Pro 80 (e56): 1–41.

Ashwell, Ken. 1991. “The Distribution of Microglia and Cell Death in the Fetal Rat Forebrain.” Developmental Brain Research 58: 1–12.

Bauer, HC, H Bauer, A Lametschwandtner, A Amberger, P Ruiz, and M Steiner. 1993. “Neovascularization and the Appearance of Morphological Characteristics of the Blood-Brain Barrier in the Embryonic Mouse Central Nervous System.” Developmental Brain Research 75 (2): 269–78.

Bayraktar, Omer Ali, Luis C Fuentealba, Arturo Alvarez-buylla, and David H Rowitch. 2015. “Astrocyte Development and Heterogeneity.” Cold Spring Harb Perspect Biol 7 (a020362): 1–16.

Berger, Daniel R, H Sebastian Seung, Jeff W Lichtman, Sean L Hill, and Marta Costa. 2018. “VAST (Volume Annotation and Segmentation Tool): Efficient Manual and Semi-Automatic Labeling of Large 3D Image Stacks.” Frontiers in Neural Circuits 12 (88): 1–15.

Bovetti, S., Y.-C. Hsieh, P. Bovolin, I. Perroteau, T. Kazunori, and A. C. Puche. 2007. “Blood Vessels Form a Scaffold for Neuroblast Migration in the Adult Olfactory Bulb.” Journal of Neuroscience 27 (22): 5976–80.

Brown, Lachlan S, Catherine G Foster, Jo-maree Courtney, Natalie E King, David W Howells, Brad A Sutherland, Johanna Jackson, and Brad A Sutherland. 2019. “Pericytes and Neurovascular Function in the Healthy and Diseased Brain.” Frontiers in Cellular Neuroscience 13 (June): 1–9.

Butovsky, Oleg, Mark P Jedrychowski, Craig S Moore, Ron Cialic, J Amanda, Galina Gabriely, Thomas Koeglsperger, et al. 2014. “Identification of a Unique TGF-β Dependent Molecular and Functional Signature in Microglia.” Nature Neuroscience 17 (1): 131–43.

Cardona, Albert, Stephan Saalfeld, Johannes Schindelin, Ignacio Arganda-carreras, Stephan Preibisch, Mark Longair, Pavel Tomancak, Volker Hartenstein, and Rodney J Douglas. 2012. “TrakEM2 Software for Neural Circuit Reconstruction.” PloS One 7 (6): 1–8.

Checchin, Daniella, Florian Sennlaub, Etienne Levavasseur, Martin Leduc, and Sylvain Chemtob. 2006. “Potential Role of Microglia in Retinal Blood Vessel Formation.” Investigative Opthalmology & Visual Science 47 (8): 3595.

Community, Blender Online. 2018. “Blender- a 3D Modelling and Rendering Package.” Blender Foundation.

Daneman, Richard. 2012. “The Blood – Brain Barrier in Health and Disease.” Annals of Neurology 72 (5): 648–72.

Daneman, Richard, Lu Zhou, Amanuel A Kebede, and Ben A Barres. 2010. “Pericytes Are Required for Blood-Brain Barrier Integrity during Embryogenesis.” Nature 468 (7323): 562– 66.

Davalos, Dimitrios, Jaime Grutzendler, Guang Yang, Jiyun V Kim, Yi Zuo, Steffen Jung, Dan R Littman, Michael L Dustin, and Wen-Biao Gan. 2005. “ATP Mediates Rapid Microglial Response to Local Brain Injury in Vivo.” Nature Neuroscience 8 (6): 752–58.

Davalos, Dimitrios, Jae Kyu Ryu, Mario Merlini, Kim M Baeten, Natacha Le Moan, Mark A Petersen, Thomas J Deerinck, et al. 2012. “Fibrinogen-Induced Perivascular Microglial Clustering Is Required for the Development of Axonal Damage in Neuroinflammation.” Nature Communications 3: 1–15.

Dudiki, Tejasvi, Julia Meller, Gautam Mahajan, Huan Liu, Irina Zhevlakova, Samantha Ste, Conner Witherow, Eugene Podrez, Chandrasekhar R Kothapalli, and Tatiana V Byzova. 2020. “Microglia Control Vascular Architecture via a TGFB1 Dependent Paracrine Mechanism Linked to Tissue Mechanics.” Nature Communications 11 (986): 1–16.

Elmore, Monica R P, Allison R. Najafi, Maya A. Koike, Nabil N. Dagher, Elizabeth E. Spangenberg, Rachel A. Rice, Masashi Kitazawa, et al. 2014. “Colony-Stimulating Factor 1 Receptor Signaling Is Necessary for Microglia Viability, Unmasking a Microglia Progenitor Cell in the Adult Brain.” Neuron 82 (2): 380–97.

Fantin, Alessandro, Joaquim M Vieira, Gaia Gestri, Laura Denti, Quenten Schwarz, Sergey Prykhozhij, Francesca Peri, Stephen W Wilson, and Christiana Ruhrberg. 2010. “Tissue Macrophages Act as Cellular Chaperones for Vascular Anastomosis Downstream of VEGF-Mediated Endothelial Tip Cell Induction.” Blood 116 (5): 829–41.

Frost, Jeffrey L., and Dorothy P. Schafer. 2016. “Microglia: Architects of the Developing Nervous System.” Trends in Cell Biology 26 (8): 587–97.

Fujioka, Teppei, Naoko Kaneko, and Kazunobu Sawamoto. 2019. “Blood Vessels as a Scaffold for Neuronal Migration.” Neurochistry International 126: 69–73.

Ginhoux, Florent, Melanie Greter, Marylene Leboeuf, Sayan Nandi, Peter See, Solen Gokhan, Mark F Mehler, et al. 2010. “Fate Mapping Analysis Reveals That Adult Microglia Derive from Primitve Macrophages.” Science (New York, N.Y.) 330 (November): 841–45.

Goldey, Glenn J., Demetris K. Roumis, Lindsey L. Glickfeld, Aaron M. Kerlin, R. Clay Reid, Vincent Bonin, Dorothy P Schafer, and Mark L. Andermann. 2014. “Versatile Cranial Window Strategies for Long-Term Two-Photon Imaging in Awake Mice.” Nature Protocols 9 (11): 2515–38.

Grant, Roger I, David A Hartmann, Robert G Underly, Narayan R Bhat, and Andy Y Shih. 2019. “Organizational Hierarchy and Structural Diversity of Microvascular Pericytes in Adult Mouse Cortex.” Journal of Cerebral Blood Flow & Metabolism 39 (3): 411–25.

Grossmann, Ruth, Nick Stence, Jenny Carr, Leah Fuller, Marc Waite, and Michael E. Dailey. 2002. “Juxtavascular Microglia Migrate along Brain Microvessels Following Activation during Early Postnatal Development.” Glia 37 (3): 229–40.

Gunner, Georgia, Lucas Cheadle, Kasey M Johnson, Pinar Ayata, Ana Badimon, Erica Mondo, M Aurel Nagy, et al. 2019. “Sensory Lesioning Induces Microglial Synapse Elimination via ADAM10 and Fractalkine Signaling.” Nature Neuroscience 22 (July): 1075–88.

Hammond, Timothy R, Daisy Robinton, and Beth Stevens. 2018. “Microglia and the Brain : Complementary Partners in Development and Disease.” Annual Review of Cell and Developmental Biology Oct 6 (34): 523–44.

Hanamsagar, Richa, and Staci D Bilbo. 2017. “Environment Matters : Microglia Function and Dysfunction in a Changing World.” Curr. Opin. Neurobiol. 47: 146–55.

Haruwaka, Koichiro, Ako Ikegami, Yoshihisa Tachibana, Nobuhiko Ohno, Hiroyuki Konishi, Akari Hashimoto, Mami Matsumoto, et al. 2019. “Dual Microglia Effects on Blood Brain Barrier Permeability Induced by Systemic Inflammation.” Nature Communications 10 (5816): 1–17.

Hoshiko, M, I Arnoux, E Avignone, N Yamamoto, and E Audinat. 2012. “Deficiency of the Microglial Receptor CX3CR1 Impairs Postnatal Functional Development of Thalamocortical Synapses in the Barrel Cortex.” J Neurosci 32 (43): 15106–11.

Kimelberg, Harold K, and Maiken Nedergaard. 2010. “Functions of Astrocytes and Their Potential As Therapeutic Targets.” Neurotherapeutics 7 (October): 338–53.

Korogod, Natalya, Carl C H Petersen, and Graham W Knott. 2015. “Ultrastructural Analysis of Adult Mouse Neocortex Comparing Aldehyde Perfusion with Cryo Fixation.” ELife 4: 1–17.

Kubota, Yoshiaki, Keiyo Takubo, Takatsune Shimizu, Hiroaki Ohno, Kazuo Kishi, Masabumi Shibuya, Hideyuki Saya, and Toshio Suda. 2009. “M-CSF Inhibition Selectively Targets Pathological Angiogenesis and Lymphangiogenesis.” The Journal of Experimental Medicine 206 (5): 1089–1102.

Kubota, Yoshiyuki, Jaerin Sohn, Sayuri Hatada, Meike Schurr, Jakob Straehle, Anjali Gour, Ralph Neujahr, Takafumi Miki, Shawn Mikula, and Yasuo Kawaguchi. 2018. “A Carbon Nanotube Tape for Serial-Section Electron Microscopy of Brain Ultrastructure.” Nature Communications 9 (437): 1–3.

Lebovitz, Yeonwoo, Veronica M Ringel-scaia, Irving C Allen, and Theus H Michelle. 2018. “Emerging Developments in Microbiome and Microglia Research : Implications for Neurodevelopmental Disorders.” Frontiers in Immunology 9 (September): 1–9.

Macvicar, Brian A, and Eric A Newman. 2015. “Astrocyte Regulation of Blood Flow in the Brain.” Cold Spring Harb Perspect Biol 7 (a020388): 1–14.

Mastorakos, Panagiotis, and Dorian Mcgavern. 2019. “The Anatomy and Immunology of Vasculature in the Central Nervous System.” Science Immunology 4: 1–14.

Mathiisen, Thomas Misje, Knut Petter Lehre, Niels Christian Danbolt, and O L E Petter Ottersen. 2010. “The Perivascular Astroglial Sheath Provides a Complete Covering of the Brain Microvessels : An Electron Microscopic 3D Reconstruction.” Glia 1103 (March): 1094–1103.

McConnell, Heather L, Cymon N Kersch, Randall L Woltjer, and Edward A Neuwelt. 2017. “The Translational Significance of the Neurovascular Unit.” The Journal of Biological Chemistry 292 (3): 762–70.

Monier, Anne, Homa Adle-Biassette, Anne-Lise Delezoide, Philippe Evrard, Pierre Gressens, and Catherine Verney. 2007. “Entry and Distribution of Microglial Cells in Human Embryonic and Fetal Cerebral Cortex.” Journal of Neuropathology and Experimental Neurology 66 (5): 372–82.

Navascués, Julio, Ruth Calvente, José L Marín-teva, and Miguel A Cuadros. 2000. “Entry, Dispersion and Differentiation of Microglia in the Developing Central Nervous System.” Anais Da Academia Brasileira de Ciencias 72 (1): 91–102.

Nikodemova, Maria, Rebecca S. Kimyon, Ishani De, Alissa L. Small, Lara S. Collier, and Jyoti J. Watters. 2015. “Microglial Numbers Attain Adult Levels after Undergoing a Rapid Decrease in Cell Number in the Third Postnatal Week.” Journal of Neuroimmunology 278: 280–88.

Nimmerjahn, Axel, Frank Kirchhoff, and Fritjof Helmchen. 2005. “Resting Microglial Cells Are Highly Dynamic Surveillants of Brain Parenchyma in Vivo.” Science (New York, N.Y.) 308 (5726): 1314–19.

Paolicelli, Rosa C, Giulia Bolasco, Francesca Pagani, Laura Maggi, Maria Scianni, Patrizia Panzanelli, Maurizio Giustetto, et al. 2011. “Synaptic Pruning by Microglia Is Necessary for Normal Brain Development.” Science (New York, N.Y.) 333 (6048): 1456–58.

Parkhurst, Christopher N., Guang Yang, Ipe Ninan, Jeffrey N. Savas, John R. Yates, Juan J. Lafaille, Barbara L. Hempstead, Dan R. Littman, and Wen Biao Gan. 2013. “Microglia Promote Learning-Dependent Synapse Formation through Brain-Derived Neurotrophic Factor.” Cell 155 (7): 1596–1609.

Parslow, Adam C, Albert Cardona, and Robert J Bryson-richardson. 2014. “Sample Drift Correction Following 4D Confocal Time-Lapse Imaging.” Journal of Visualized Experiments : JoVE 86 (e51086).

Perry, V.H., D.A. Hume, and S. Gordon. 1985. “Immunohistochemical Localization of Macrophages and Microglia in the Adult and Developing Mouse Brain.” Neuroscience 15 (2): 313–26.

Rothhammer, Veit, Davis M Borucki, Emily C Tjon, Maisa C Takenaka, Alberto Ardura Fabregat, Kalil Alves De Lima, Cristina Gutierrez Vazquez, et al. 2018. “Microglial Control of Astrocytes in Response to Microbial Metabolites.” Nature 557 (7707): 724–28.

Rymo, Simin F., Holger Gerhardt, Fredrik Wolfhagen Sand, Richard Lang, Anne Uv, and Christer Betsholtz. 2011. “A Two-Way Communication between Microglial Cells and Angiogenic Sprouts Regulates Angiogenesis in Aortic Ring Cultures.” PLoS ONE 6 (1).

Saili, Katerine S, Todd J Zurlinden, Andrew J Schwab, Aymeric Silvin, C Nancy, E Sidney Hunter Iii, Florent Ginhoux, and Thomas B Knudsen. 2017. “Blood-Brain Barrier Development: Systems Modeling and Predictive Toxicology.” Birth Defects Research 109 (20): 1680–1710.

Schafer, Dorothy P, Emily K Lehrman, Amanda G Kautzman, Ryuta Koyama, Alan R Mardinly, Ryo Yamasaki, Richard M Ransohoff, Michael E Greenberg, Ben A Barres, and Beth Stevens. 2012. “Microglia Sculpt Postnatal Neural Circuits in an Activity and Complement-Dependent Manner.” Neuron 74 (4): 691–705.

Schiweck, Juliane, Britta J Eickholt, and Kai Murk. 2018. “Important Shapeshifter : Mechanisms Allowing Astrocytes to Respond to the Changing Nervous System During Development, Injury and Disease.” Frontiers in Cellular Neuroscience 12 (261): 1–17.

Smolders, Sophie Marie Thérèse, Nina Swinnen, Sofie Kessels, Kaline Arnauts, Silke Smolders, Barbara Le Bras, Jean Michel Rigo, Pascal Legendre, and Bert Brône. 2017. “Age-Specific Function of Α5β1 Integrin in Microglial Migration during Early Colonization of the Developing Mouse Cortex.” Glia 65: 1072–88.

Sorokin, Sergei P, Richar F Hoyt, Dana G Blunt, and Nancy A Mcnellyl. 1992. “Macrophage Develoment: II. Early Ontogeny of Macrophage Pupulations in the Brain, Liver, and Lungs of Rat Embryos as Revealed by a Lectin Marker.” The Anatomical Record 232 (4): 527–50.

Stankovic, Nevenka Dudvarski, Marcin Teodorczyk, and Robert Ploen. 2016. “Microglia – Blood Vessel Interactions : A Double-Edged Sword in Brain Pathologies.” Acta Neuropathologica 131 (3): 347–63.

Swinnen, Nina, Sophie Smolders, Ariel Avila, Kristof Notelaers, Rik Paesen, Marcel Ameloot, Bert Brône, Pascal Legendre, and Jean Michel Rigo. 2013. “Complex Invasion Pattern of the Cerebral Cortex by Microglial Cells during Development of the Mouse Embryo.” Glia 61 (2): 150–63.

Tapia, Juan C, Narayanan Kasthuri, Kenneth Hayworth, Richard Schalek, Jeff W Lichtman, Stephen J Smith, and Joann Buchanan. 2013. “High Contrast En Bloc Staining of Neuronal Tissue for Field Emission Scanning Electron Microscopy.” Nature Protocols 7 (2): 193–206.

Tinevez, Jean-yves, Nick Perry, Johannes Schindelin, Genevieve M Hoopes, Gregory D Reynolds, Emmanuel Laplantine, Sebastian Y Bednarek, Spencer L Shorte, and Kevin W Eliceiri. 2017. “TrackMate : An Open and Extensible Platform for Single-Particle Tracking.” Methods 115: 80–90.

Tremblay, Marie-Eve, Rebecca L. Lowery, and Ania K. Majewska. 2010. “Microglial Interactions with Synapses Are Modulated by Visual Experience.” PLoS Biology 8 (11).

Tsai, H.-H., J. Niu, R. Munji, D. Davalos, J. Chang, H. Zhang, A.-C. Tien, et al. 2016. “Oligodendrocyte Precursors Migrate along Vasculature in the Developing Nervous System.” Science (New York, N.Y.) 351 (6271): 379–84.

Ueno, Masaki, and Toshihide Yamashita. 2014. “ScienceDirect Bidirectional Tuning of Microglia in the Developing Brain : From Neurogenesis to Neural Circuit Formation.” Current Opinion in Neurobiology 27: 8–15.

Welker, Carol, and Thomas A Woolsey. 1974. “Structure of Layer IV in the Somatosensory Neocortex of the Rat: Description and Comparison with the Mouse.” Journal of Comparative Neurology 158: 437–53.

Whitman, Mary C, Wen Fan, Lorena Rela, Diego J Rodriguez-gil, and Charles A Greer. 2009. “Blood Vessels Form a Migratory Scaffold in the Rostral Migratory Stream.” Journal of Comparative Neurology 516 (2): 94–104.

Woolsey, Thomas A., and Hendrik Van der Loos. 1970. “The Structural Organization of Layer IV in the Somatosensory Region (SI) of Mouse Cerebral Cortex.” Brain Research 17: 205–42.

Yamanishi, Emiko, Masanori Takahashi, Yumiko Saga, and Noriko Osumi. 2012. “Penetration and Differentiation of Cephalic Neural Crest-Derived Cells in the Developing Mouse Telencephalon.” Development, Growth, and Differentiation 54: 785–800.

Zeisel, Amit, Ana B Munoz-Monchado, Simone Codeluppi, Peter Lonnerberg, Gioele La Manno, Anna Jureus, Sueli Marques, et al. 2015. “Cell Types in the Moues Cortex and Hippocampus Revealed by Single-Cell RNA-Seq.” Science (New York, N.Y.) 347 (6226): 1138–43.

Zhan, Yang, Rosa C Paolicelli, Francesco Sforazzini, Laetitia Weinhard, Giulia Bolasco, Francesca Pagani, Alexei L Vyssotski, et al. 2014. “Deficient Neuron-Microglia Signaling Results in Impaired Functional Brain Connectivity and Social Behavior.” Nature Neuroscience 17 (3): 400–406.

Zhao, Xiaoliang, Ukpong B Eyo, Madhuvika Murugan, and Long-jun Wu. 2018. “Microglial Interactions with the Neurovascular System in Physiology and Pathology.” Developmental Neurobiology 78 (6): 604–17.

Zlokovic, Berislav V. 2008. “The Blood-Brain Barrier in Health and Chronic Neurodegenerative Disorders.” Neuron 57: 178–201.

